# Photo sensing and quorum sensing are integrated to control bacterial group behaviors

**DOI:** 10.1101/747618

**Authors:** Sampriti Mukherjee, Matthew Jemielita, Vasiliki Stergioula, Mikhail Tikhonov, Bonnie L. Bassler

**Affiliations:** Princeton University, Department of Molecular Biology, Princeton, NJ 08544, USA; Physics department, Washington University in St Louis, St Louis, MO 63130, USA; Howard Hughes Medical Institute, Chevy Chase, MD 20815, USA

**Keywords:** bacteria, *Pseudomonas*, quorum sensing, photo sensing, biofilms, virulence, two-component system

## Abstract

*Pseudomonas aeruginosa* transitions between the free-swimming state and the sessile biofilm mode during its pathogenic lifestyle. We show that quorum sensing represses *P. aeruginosa* biofilm formation and virulence by activating expression of genes encoding the KinB-AlgB two-component system. Phospho-AlgB represses biofilm and virulence genes, while KinB dephosphorylates, and thereby, inactivates AlgB. We discover that the photoreceptor BphP is the kinase that, in response to light, phosphorylates and activates AlgB. Indeed, exposing *P. aeruginosa* to light represses biofilm formation and virulence gene expression. To our knowledge, *P. aeruginosa* was not previously known to detect light. The KinB-AlgB-BphP module is present in all Pseudomonads, and we demonstrate that AlgB is the cognate response regulator for BphP in diverse bacterial phyla. We propose that KinB-AlgB-BphP constitutes a “three-component” system and AlgB is the node at which varied sensory information is integrated. This study sets the stage for light-mediated control of *P. aeruginosa* infectivity.

## INTRODUCTION

Bacterial responses to self-generated and exogenous stimuli influence their survival, persistence in particular niches, and lifestyle transitions, such as alterations between being free-swimming or existing as a member of a biofilm. Biofilms are three-dimensional structured communities of bacterial cells encased in an extracellular matrix (Flemming and Wingender, 2010; Flemming et al., 2016). Bacteria living in biofilms exhibit superior resilience to environmental stresses such as antimicrobials and host immune responses (Flemming et al., 2016; Koo et al., 2017). While many cues are known to drive the biofilm-planktonic transition, it is largely mysterious how sensory information is detected, integrated, and transduced to control alterations between the two lifestyles. Here, we show that photo sensing and quorum sensing converge to control biofilm formation and virulence in the global pathogen *Pseudomonas aeruginosa*, and we define the pathway connecting the light and quorum sensing inputs to the virulence and biofilm outputs.

Light is a common environmental cue that is detected by photoreceptors present in all domains of life (Horst et al., 2007; Kottke et al., 2018). Particular photoreceptor photosensory domains are activated by specific wavelengths of light (Shcherbakova et al., 2015). In bacteria, the most abundant photoreceptors are phytochromes (Gomelsky and Hoff, 2011), typically possessing an amino-terminal chromophore-binding domain and a carboxy-terminal histidine kinase (HK) domain. Bacteriophytochromes assemble with the chromophore called biliverdin (Gourinchas et al., 2019). Surprisingly, very few bacteria encode a cognate response regulator (RR) in close proximity to the gene specifying the bacteriophytochrome (Beattie et al., 2018), leaving the systems mostly undefined.

Another extracellular parameter monitored by bacteria is their cell-population density. To do this, bacteria use the cell-to-cell communication process called quorum sensing, which relies on production and detection of extracellular signaling molecules called autoinducers (Mukherjee and Bassler, 2019). Quorum sensing allows groups of bacteria to synchronously alter behavior in response to changes in the population density and species composition of the vicinal community. Many pathogenic bacteria require quorum sensing to establish successful infections (Rutherford and Bassler, 2012).

In the human pathogen *Pseudomonas aeruginosa*, quorum sensing is required for virulence and biofilm formation (Davies et al., 1998; Mukherjee et al., 2017; Rumbaugh et al., 2000). In this study, we examine the mechanism by which the *P. aeruginosa* RhlR quorum-sensing receptor represses biofilm formation. A genetic screen reveals that RhlR activates the expression of the *algB-kinB* operon encoding a two-component system (TCS) in which KinB and AlgB are the sensor HK and cognate RR, respectively (Figure 1). We find that AlgB∼P is a repressor of biofilm formation and virulence gene expression. KinB is a phosphatase that dephosphorylates, and thereby inactivates AlgB. Using genetic suppressor analysis, we discover that BphP is the HK that phosphorylates and activates AlgB, enabling AlgB to repress biofilm formation and genes encoding virulence factors (Figure 1). BphP is a far-red light sensing bacteriophytochrome (Tasler et al., 2005), and indeed, we demonstrate that *P. aeruginosa* biofilm formation and virulence gene expression are repressed by far-red light. Phylogenetic analyses show that the KinB-AlgB-BphP module is conserved in all Pseudomonads and, moreover, AlgB is present in the majority of bacteria that possess BphP orthologs. This final finding suggests that the BphP-AlgB interaction is widespread. As proof of this notion, we show that *P. aeruginosa* BphP can phosphorylate AlgB orthologs from α-, β-, and γ-Proteobacteria. Thus, KinB-AlgB-BphP constitute a “three-component” system, and we propose that AlgB functions as the integrator that conveys multiple environmental cues including those specifying population density and the presence or absence of light into the regulation of collective behaviors (Figure 1). We further predict that AlgB functions as the cognate RR for BphP in all bacteria that possess BphP as an orphan HK. The downstream signal transduction components and the outputs of photo sensory cascades are not known in the majority of non-photosynthetic bacteria that possess them making their physiological roles difficult to discern. This study provides the entire cascade—light as the input, BphP as the detector, AlgB as the signal transducer, and biofilm formation and virulence factor production as the outputs—enabling unprecedented insight into light-driven control of bacterial behavior.

**Figure 1:**
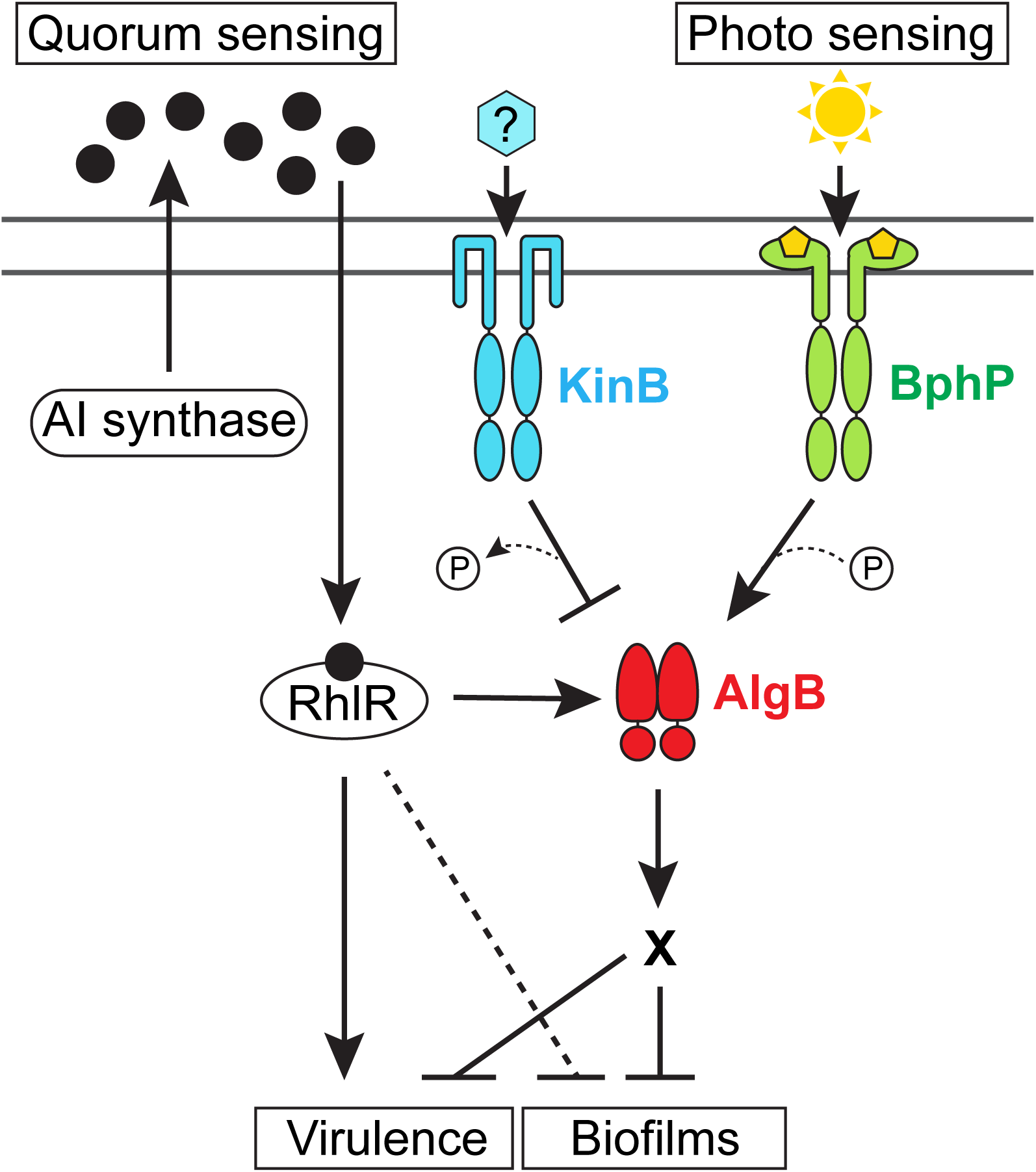
Model for *P. aeruginosa* integration of quorum-sensing and photo-sensing information into the control of virulence and biofilm development. The RhlR quorum-sensing receptor binds its cognate autoinducer (AI) produced by either the RhlI or PqsE autoinducer synthase (black circles) at high cell density (Mukherjee et al., 2018). The RhlR-AI complex represses biofilm formation and virulence gene expression by activating transcription of the *algB-kinB* operon encoding the KinB HK and the AlgB RR, the latter a repressor of biofilm formation. KinB antagonizes AlgB by dephosphorylation. The stimulus (blue hexagon) for KinB is unknown. Photo sensing stimulates the BphP HK to auto-phosphorylate, and subsequently transfer the phosphoryl group to AlgB to activate AlgB. AlgB∼P activates transcription of genes required for repression of group behaviors such as biofilm formation and virulence. A “P” in a circle denotes addition or removal of a phosphate moiety. X denotes that the genes functioning downstream of AlgB in the process are not known. The RhlR-AI complex directly activates virulence gene expression and also represses biofilm formation by additional unknown mechanisms (dotted line).

## RESULTS

### KinB activates and AlgB represses RhlR-dependent group behaviors

We recently discovered that the *P. aeruginosa* quorum-sensing receptor RhlR represses biofilm formation (Mukherjee et al., 2017, 2018). Specifically, on Congo red agar biofilm medium, wildtype (WT) *P*. *aeruginosa* UCBPP-PA14 (hereafter called PA14) exhibits a rugose-center/smooth-periphery colony biofilm phenotype, while the Δ*rhlR* mutant forms a larger hyper-rugose biofilm (Figure 2A). To determine the mechanism by which RhlR impedes biofilm formation, we randomly mutagenized the Δ*rhlR* strain using the Tn*5* IS50L derivative IS*lacZ*/*hah* (Jacobs et al., 2003) and screened for colonies exhibiting either a WT or a smooth colony biofilm phenotype. Our rationale was that inactivation of a gene(s) encoding a component that functions downstream of RhlR in biofilm formation would sever the connection between RhlR and repression of biofilm formation. We screened 5,000 transposon insertion mutants. Strains harboring insertions located in genes encoding hypothetical proteins, proteins involved in twitching motility, and proteins required for Pel polysaccharide synthesis all produced smooth colony biofilms (Table S1). Most of these genes were already known to play roles in *P. aeruginosa* biofilm formation (Fazli et al., 2014). Here, we focus on one transposon insertion mutant that exhibited a smooth colony biofilm phenotype that mapped to the gene *PA14_72390* encoding the KinB transmembrane HK (Figure 2A) (Chand et al., 2011). *kinB* is located immediately downstream of *algB* in a di-cistron that is conserved in all sequenced Pseudomonads (Figure S1). To verify that KinB plays a role in biofilm formation, we generated an in-frame marker-less deletion of *kinB* in the chromosomes of the WT and the Δ*rhlR* strains. Both the Δ*kinB* single and Δ*rhlR* Δ*kinB* double mutants failed to form biofilms and instead exhibited smooth colony phenotypes (Figure 2A). Introduction of a plasmid carrying the *kinB* gene conferred a hyper-rugose phenotype to the WT and restored biofilm formation to the Δ*kinB* and Δ*rhlR* Δ*kinB* mutants (Figure 2A). By contrast, introduction of a plasmid carrying *rhlR* did not alter the smooth biofilm phenotype of the Δ*rhlR* Δ*kinB* mutant (Figure 2A). We conclude that, in *P. aeruginosa*, KinB is essential for biofilm formation, KinB is an activator of biofilm formation, and KinB functions downstream of RhlR in the biofilm formation process.

**Figure 2:**
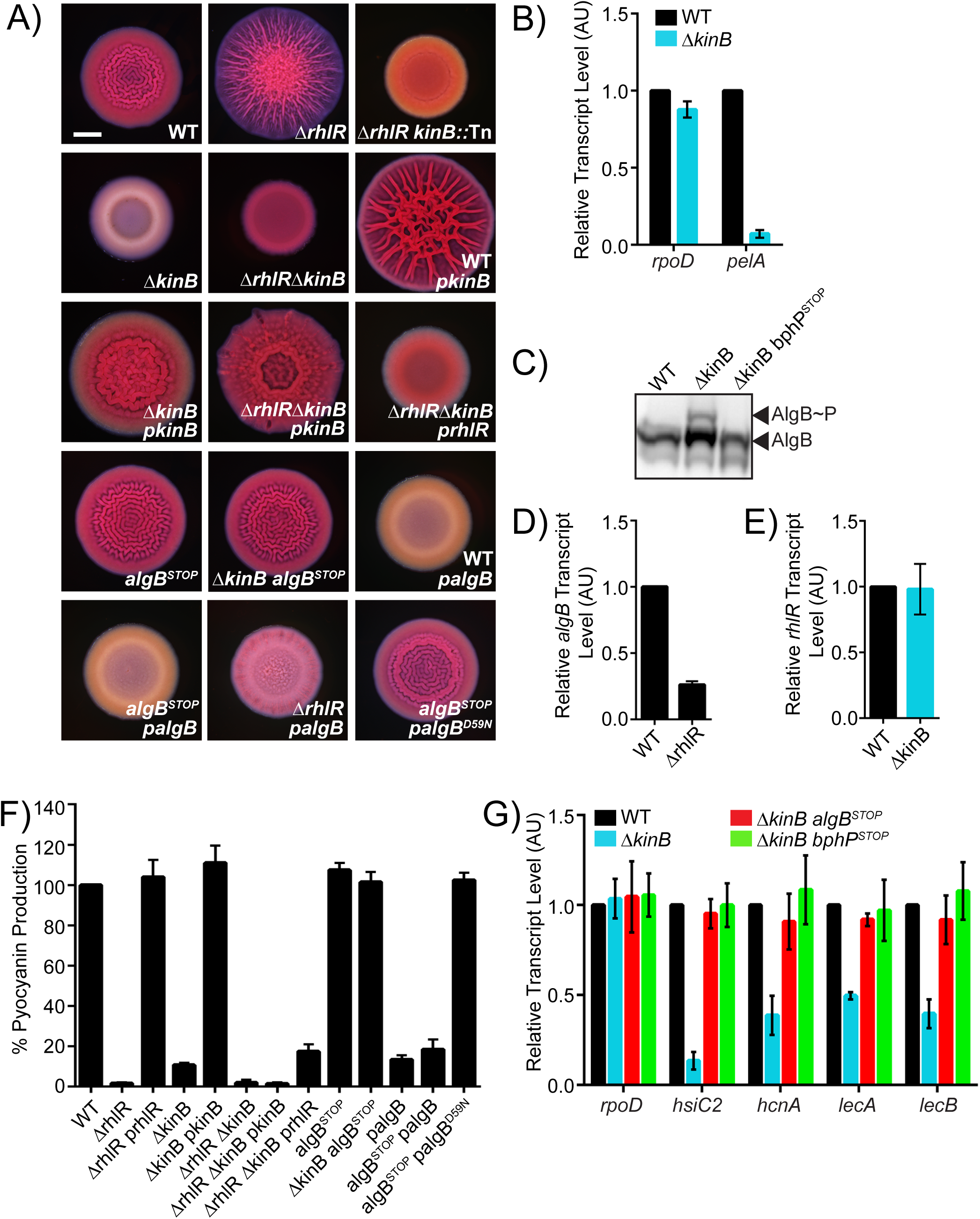
RhlR represses biofilm formation via KinB. A) Colony biofilm phenotypes of WT PA14 and the designated mutants on Congo red agar medium after 72 h of growth. *kinB*::Tn refers to a mutant identified in a genetic screen harboring a transposon insertion in *kinB*. *pkinB, prhlR*, and *palgB* refer to *kinB, rhlR*, and *algB*, respectively, under the P_lac_ promoter on pUCP18. Scale bar for all images is 2 mm. B) Relative expression levels of *rpoD* and *pelA* measured by qRT-PCR in WT and Δ*kinB* mutant biofilms grown as in (A). C) Phos-tag Western blot analysis of the indicated strains probed for 3xFLAG-AlgB. D) Relative *algB* transcript levels measured by qRT-PCR in WT PA14 and the Δ*rhlR* mutant grown planktonically to OD_600_ = 1.0. E) Relative *rhlR* transcript levels measured by qRT-PCR in WT PA14 and the Δ*kinB* mutant grown planktonically to OD_600_ = 1.0. F) Pyocyanin production (OD_695_) was measured in WT PA14 and the designated mutants. Production from the WT was set to 100%. G) Relative expression of *rpoD*, *hsiC2*, *hcnA*, *lecA*, and *lecB* measured by qRT-PCR in WT PA14 and the designated mutants grown planktonically to OD_600_ = 1.0. *rpoD* is used as the control for comparison. For panels B, D, E and G, data were normalized to 16S RNA levels and the WT levels were set to 1.0. AU denotes arbitrary units. For data in panels B, D, E, F, and G, error bars represent standard error of the mean (SEM) for three biological replicates.

PA14 requires Pel, the primary biofilm matrix exopolysaccharide for biofilm formation (Friedman and Kolter, 2004) (Note: PA14 does not produce the Psl exopolysaccharide and alginate does not contribute significantly to the PA14 biofilm matrix, unlike in *P. aeruginosa* PAO1 (Wozniak et al., 2003)). To examine if the mechanism by which KinB alters biofilm formation is by changing Pel production, we performed quantitative RT-PCR analyses on WT and Δ*kinB* biofilms probing for the expression of the housekeeping gene *rpoD* and the Pel biosynthetic gene *pelA* (Figure 2B). Expression of *rpoD* did not change between the WT and the Δ*kinB* mutant, while transcription of *pelA* was ∼14-fold lower in the Δ*kinB* strain than in the WT. We conclude that KinB activates Pel production, which is why KinB is required for PA14 biofilm formation.

KinB is a transmembrane HK that undergoes autophosphorylation and then transfers the phosphate to its cognate RR AlgB (Ma et al., 1997). To determine if AlgB functions downstream of KinB to control biofilm formation, we engineered a stop codon in the *algB* gene to obtain an *algB^STOP^* mutant. This strategy enabled us to prevent AlgB translation without affecting transcription of *kinB*. The *algB^STOP^* mutant had a biofilm phenotype indistinguishable from the WT (Figure 2A). However, introduction of the *algB^STOP^* mutation into the Δ*kinB* strain restored biofilm formation (Figure 2A). Furthermore, overexpression of *algB* repressed biofilm formation in the WT as evidenced by the resulting smooth colony biofilm phenotype (Figure 2A). Overexpression of *algB* also repressed biofilm formation in the *algB^STOP^* and Δ*rhlR* strains (Figure 2A). Thus, KinB activates while AlgB represses biofilm development.

AlgB has an amino-terminal domain containing the site of phosphorylation (residue D59), a central ATP-binding domain, and a carboxy-terminal helix-turn-helix motif for binding DNA (Figure S2) (Ma et al., 1998). AlgB is a member of the NtrC subfamily of RRs and it possesses the hallmark GAFTGA motif required for interaction with RpoN (σ^54^) (Wozniak and Ohman, 1991). Typically, NtrC-type RRs act as transcriptional activators when they are phosphorylated (reviewed in Bush and Dixon, 2012). To investigate if phosphorylation of AlgB is required for repression of biofilm formation, we substituted the aspartate at residue 59 with an asparagine residue to preclude phosphorylation. We overexpressed the *algB^D59N^* allele in the PA14 strain carrying the *algB^STOP^* mutation. Unlike WT AlgB, AlgB^D59N^ failed to repress biofilm formation (Figure 2A). To ensure the validity of this result, we generated amino-terminal 3xFLAG tagged *algB* and *algB^D59N^* fusions and expressed them from a plasmid in the *algB^STOP^* mutant. Western blot showed that both proteins are stable (Figure S3A). We conclude that the phosphorylated form of AlgB is active and is required for AlgB-mediated repression of biofilm development. We presume that AlgB∼P functions indirectly as a transcriptional activator to promote the expression of a gene(s) encoding a negative regulator of biofilm formation (Figure 1).

Our results show that AlgB functions downstream of KinB and that KinB and AlgB have opposing activities with respect to PA14 biofilm formation. *In vitro*, KinB possess both kinase and phosphatase activities (Chand et al., 2012). One mechanism by which KinB could antagonize AlgB function is by acting as a phosphatase that dephosphorylates AlgB, rendering it inactive. To test this possibility, we integrated the 3xFLAG tagged *algB* allele at the native *algB* locus in the chromosomes of WT PA14 and the Δ*kinB* mutant. Biofilm analyses show that 3xFLAG-AlgB is functional (Figure S3B). Next, we assessed the phosphorylation status of 3xFLAG-AlgB *in vivo*. Figure 2C shows that AlgB∼P accumulates in the Δ*kinB* mutant compared to in the WT. To verify these claims regarding the signal transduction mechanism, we engineered a missense mutation into KinB at a conserved proline (P390) that is required for phosphatase activity (Chand et al., 2012). Specifically, we generated both *kinB-SNAP* and *kinB^P390S^-SNAP* fusions and introduced these alleles at the native *kinB* locus on the chromosome of *P. aeruginosa*. Carboxy-terminal tagging of KinB with SNAP does not interfere with its function as the strain carrying *kinB-SNAP* forms biofilms that are indistinguishable from those of WT PA14 (Figure S3C and D). The KinB^P390S^-SNAP protein is also produced and stable (Figure S3C), however, identical to the Δ*kinB* mutant, the strain carrying *kinB^P390S^-SNAP* fails to form biofilms (Figure S3D). These data demonstrate that KinB acts as a phosphatase to inhibit AlgB function *in vivo*. We therefore hypothesize, and we come back to this point below, that some other HK must phosphorylate AlgB to activate it and enable it to function as a repressor of biofilm development.

Our data show that the KinB-AlgB TCS functions downstream of RhlR to repress biofilm formation. An obvious mechanism by which RhlR could influence KinB-AlgB activity is by activating transcription of the *algB-kinB* operon. Indeed, RT-PCR shows that *algB-kinB* transcript levels are ∼4-fold higher in the WT than in the Δ*rhlR* mutant (Figure 2D). Thus, RhlR activates expression of *algB-kinB* operon. By contrast, deletion of *kinB* has no effect on *rhlR* transcript levels (Figure 2E), confirming their epistatic relationship.

KinB has been reported to be required for pyocyanin production (Chand et al., 2011). Pyocyanin is a RhlR-dependent virulence factor (Brint and Ohman, 1995; Mukherjee et al., 2017). Our findings of a regulatory connection between KinB and RhlR suggest that KinB and RhlR could jointly regulate pyocyanin production. To test this idea, we measured pyocyanin production in planktonic cultures of WT, Δ*rhlR*, Δ*kinB*, and Δ*rhlR* Δ*kinB* strains. Similar to what has been reported previously, deletion of *rhlR* and/or *kinB* abolished pyocyanin production (Figure 2F). Overexpression of *rhlR* in the Δ*rhlR* strain and overproduction of *kinB* in the Δ*kinB* strain restored pyocyanin production, demonstrating that our expression constructs are functional (Figure 2F). By contrast, overexpression of either *rhlR* or *kinB* in the Δ*rhlR* Δ*kinB* double mutant failed to rescue pyocyanin production (Figure 2F). Thus, RhlR and KinB are both required activators of pyocyanin production in PA14. Consistent with AlgB functioning as the RR for KinB, inactivation of AlgB (i.e., *algB^STOP^*) in the Δ*kinB* background restored WT levels of pyocyanin production while overexpression of AlgB in the WT and the *algB^STOP^* mutant reduced pyocyanin levels (Figure 2F). Lastly, unlike WT AlgB, overexpression of AlgB^D59N^ failed to repress pyocyanin production suggesting that phosphorylation of AlgB is required for AlgB activity (Figure 2F).

To further explore the role of the KinB-AlgB TCS on RhlR-driven gene expression, we quantified expression of four other RhlR-activated genes (Mukherjee et al., 2017): *hsiC2* (type-VI secretion), *hcnA* (hydrogen cyanide synthase), *lecA* (galactose-binding lectin), and *lecB* (fucose-binding lectin), all encoding virulence factors, in the WT and the Δ*kinB* mutant. Expression of all four genes was lower in the Δ*kinB* mutant than in the WT (Figure 2G). Introduction of the *algB^STOP^* mutation into the Δ*kinB* mutant restored expression of all four virulence genes to WT levels (Figure 2G). Thus, both RhlR and KinB activate virulence gene expression in *P. aeruginosa*. Moreover, we conclude that AlgB is epistatic to KinB for all the phenotypes tested here, and thus KinB and AlgB function in the same pathway, albeit in opposing manners, to control biofilm formation and virulence factor production.

### The bacteriophytochrome BphP is the HK required to activate AlgB to mediate repression of quorum-sensing-controlled behaviors

We have invoked the existence of a putative HK to activate AlgB via phosphorylation. To identify this component, we used genetic suppressor analysis reasoning that mutants with defects in the upstream component required to phosphorylate AlgB would render AlgB non-functional. We further reasoned that such suppressor mutants would transform the Δ*kinB* smooth colony biofilm phenotype back to the rugose phenotype because in such mutants, AlgB could not act as a repressor of biofilm formation. We isolated 12 spontaneously-arising rugose mutants from Δ*kinB* smooth colony biofilms and analyzed them by whole genome sequencing (Figure 3A). Eight suppressors contained deletions or missense mutations in the *algB* gene, while the remaining four suppressors harbored mutations in the *bphP* gene (Figure 3B, Table S2). *bphP* is located in a di-cistron immediately downstream of *bphO* (Figure 3B, S1). We discuss *bphO* below; here we focus on *bphP*. Exactly analogous to mutation of *algB*, mutation of *bphP* was epistatic to *kinB* for all of the phenotypes tested. Specifically, engineering a STOP codon into the *bphP* gene showed no effect in WT PA14, but it restored biofilm formation, pyocyanin production, and virulence gene expression to the Δ*kinB* mutant (Figures 3C, D, and 2G). Consistent with BphP being required to activate AlgB, unlike in the WT, in the *bphP^STOP^* mutant, overexpression of *algB* failed to repress biofilm formation and pyocyanin production (Figure 3C, D). Furthermore, while overexpression of *bphP* in the WT reduced pyocyanin production to the levels of the Δ*kinB* mutant, overexpression of *bphP* had no effect in the *algB^STOP^* mutant (Figure 3D). There is a severe growth defect associated with the overexpression of *bphP*. For this reason, in Figure 3D, rather than using plasmid pUCP18, we expressed *bphP* from the low copy number plasmid pBBR1-MCS5. Unfortunately, the presence of the empty pBBR-MCS5 plasmid in WT and mutant PA14 strains abrogates biofilm formation, so we could not perform the companion biofilm assay to test overexpression of *bphP*. Nonetheless, we can conclude from Figure 3C and 3D that BphP is necessary and sufficient to activate AlgB.

**Figure 3:**
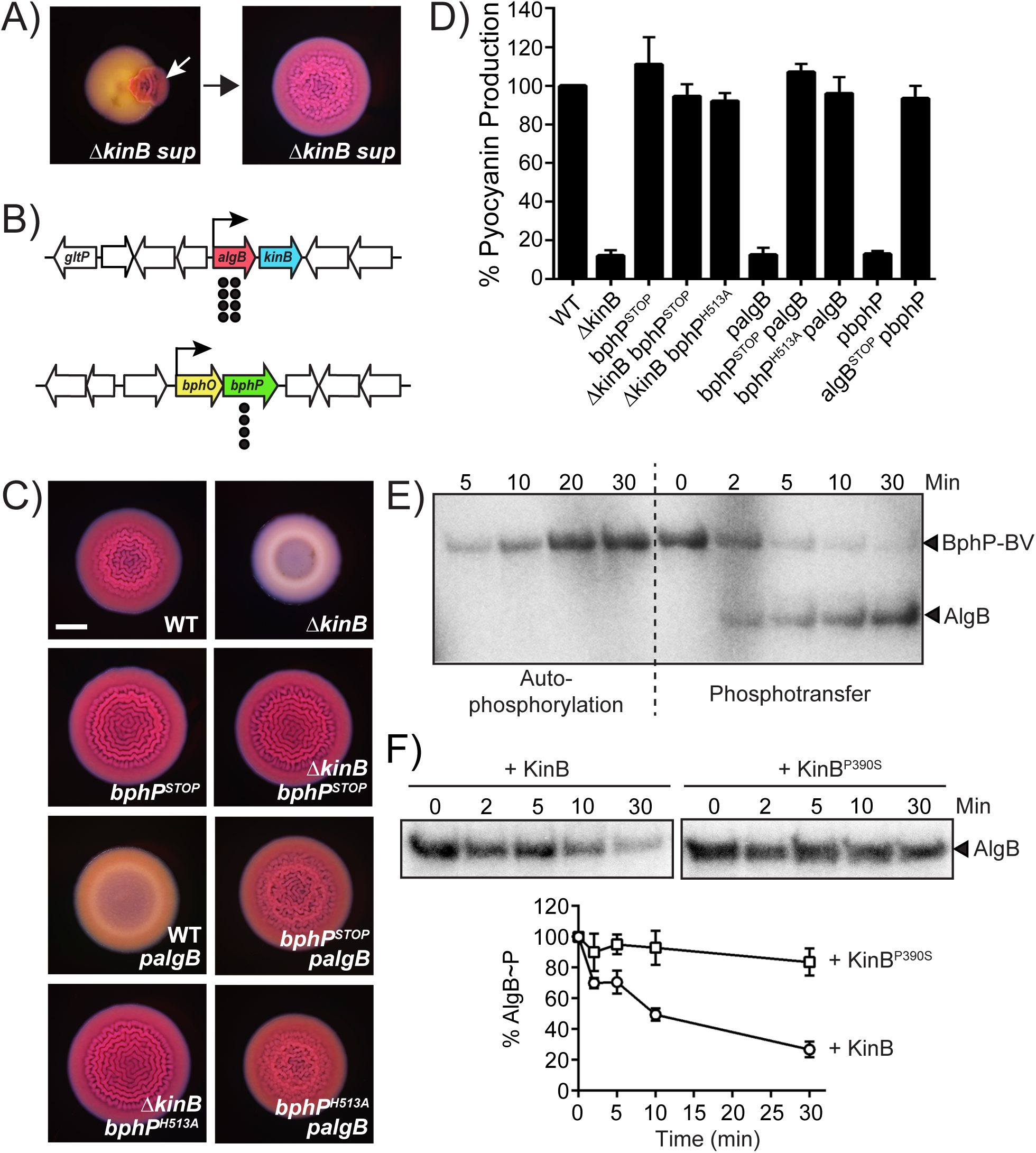
BphP is the cognate HK for AlgB. A) Shown is a representative isolation of a suppressor mutation of the Δ*kinB* smooth biofilm phenotype. The white arrow in the left panel indicates a region of rugose sectoring in the Δ*kinB* smooth biofilm that is diagnostic of the emergence of a suppressor mutation. The right panel shows the biofilm phenotype of a mutant following isolation. B) Chromosomal arrangements of the *algB* (red), *kinB* (blue), *bphO* (yellow), and *bphP* (green) genes. Large white arrows represent open reading frames (lengths not to scale), black bent arrows indicate promoters, and black circles indicate the locations of suppressor mutations. C) Colony biofilm phenotypes of WT PA14 and the designated mutants on Congo red agar medium after 72 h of growth. *palgB* refers to *algB* under the P_lac_ promoter on the pUCP18 plasmid. Scale bar is 2 mm for all images. D) Pyocyanin production (OD_695_) was measured in WT PA14 and the designated mutants. *pbphP* refers to *bphP* under the P_lac_ promoter on the pBBR-MCS5 plasmid. Error bars represent SEM for three biological replicates. E) Autophosphorylation of BphP-BV and phosphotransfer to AlgB. (Left) Autophosphorylation of BphP-BV was carried out for 30 min and samples were removed at the indicated times for electrophoresis. (Right) An equimolar amount of AlgB was added to P∼BphP-BV for 30 min and samples were removed at the indicated times for electrophoresis. F) Dephosphorylation of AlgB∼P by KinB or KinB^P390S^. Phosphotransfer to AlgB from P∼BphP-BV was carried out for 30 min. ATP was removed from the reaction, and either KinB or KinB^P390S^ was added. Samples were removed at the indicated times for electrophoresis. The top panel shows representative images of gels. The bottom graph shows % AlgB∼P levels at each time point with SEM for three independent replicates. Band intensities for AlgB∼P when KinB was added (circles) and when KinB^P390S^ was added (squares) were normalized to the level at time zero level.

BphP is a bacteriophytochrome that assembles with its chromophore biliverdin, which is produced by the heme oxygenase BphO (Figure 3B and S1) to generate a photo-sensing HK that is activated by light (Bhoo et al., 2001). *P. aeruginosa* BphP contains the HDLRNPL motif that often contains the histidine residue that undergoes autophosphorylation in transmembrane HKs (Bhate et al., 2015). In *P. aeruginosa* BphP, this histidine is residue 513. To determine if BphP kinase activity is required for AlgB activation, we generated the *bphP^H513A^* mutation, fused it to 3x*FLAG*, and introduced it onto the chromosome of the Δ*kinB* mutant. The BphP^H513A^-3xFLAG protein is produced and stable (Figure S3E), and identically to the *bphP^STOP^* allele, the *bphP^H513A^* mutation restored biofilm formation and pyocyanin production to the Δ*kinB* mutant (Figure 3C, D). Moreover, overexpression of *algB* in the *bphP^H513A^* mutant failed to repress biofilm formation and pyocyanin production (Figure 3C, D). These results show that BphP H513 and AlgB D59 are required for signal transmission, and the signal is presumably phosphorylation.

To assess phospho-relay between BphP and AlgB, we used our 3xFLAG-AlgB *in vivo* construct. In addition to introducing it into the chromosome of WT PA14, we engineered it onto the chromosome of the *bphP^STOP^* mutant. Consistent with BphP being the kinase for AlgB, Figure 2C shows that the Δ*kinB bphP^STOP^* mutant lacks the band corresponding to AlgB∼P. These data suggest that BphP transfers phosphate to AlgB. To verify this finding, we performed *in vitro* phospho-transfer assays. We purified recombinant BphP and formed a complex with it and commercially-available biliverdin (BV) to obtain the BphP-BV chromoprotein. Upon incubation with radiolabeled ATP, BphP-BV underwent autophosphorylation (Figure 3E). BphP-BV readily transferred radiolabeled phosphate to purified AlgB but not to AlgB^D59N^ (Figures 3E, S4A). Purified BphP^H513A^ complexed with BV failed to autophosphorylate and thus could not transfer phosphate to AlgB (Figure S4A). Together, these data show that BphP-BV phosphorylates and thereby activates AlgB.

Our data suggest that KinB dephosphorylates AlgB while BphP phosphorylates AlgB. To directly test this hypothesis, we reconstituted the BphP-AlgB-KinB phosphorelay *in vitro*. We purified the recombinant KinB and KinB^P390S^ proteins and added them separately, at equimolar concentration, to AlgB∼P pre-phosphorylated by BphP-BV. Figures 3F shows that over time, KinB dephosphorylates AlgB while AlgB∼P levels remain unchanged in the presence of KinB^P390S^. As control experiments, we added either KinB or KinB^P390S^ to AlgB in the presence of ATP but in the absence of BphP-BV. Both KinB and KinB^P390S^ underwent autophosphorylation and transferred phosphate to AlgB *in vitro*, but only WT KinB acted as a phosphatase to dephosphorylate AlgB (Figure S4B-D). Although our findings show that KinB is a dual kinase/phosphatase, under our *in vivo* conditions, only KinB phosphatase activity was detected. Perhaps KinB can function as a kinase for AlgB when its stimulus is present (Figure 1). Identifying the natural signal that drives the KinB kinase activity is the subject of our future work. We conclude that BphP-AlgB-KinB forms a “three-component” system in which the RR AlgB is activated by the kinase activity of the HK BphP and inhibited by the phosphatase activity of the HK KinB.

### BphP-mediated photo sensing represses *P. aeruginosa* quorum-sensing-controlled behaviors

The *P. aeruginosa* BphP bacteriophytochrome has been studied *in vitro* and its kinase activity is reported to be activated by light (Bhoo et al., 2001). To explore whether BphP photo sensing has any effect on AlgB-controlled group behaviors *in vivo*, we compared biofilm formation by WT, Δ*kinB*, Δ*kinB bphP^STOP^*, and Δ*kinB algB^STOP^* PA14 strains in the dark and under different light conditions. We note that all of the biofilm experiments in the previous sections were performed under ambient light. First, we consider WT PA14 and the Δ*kinB* mutant in the no light condition. Figure 4A shows that, in the dark, both strains formed biofilms that were indistinguishable from one another. We interpret these results to mean that in the absence of light, the BphP kinase is inactive in both WT PA14 and the Δ*kinB* mutant, AlgB is not phosphorylated, so it too is inactive, and thus, no repression of biofilm formation occurs (Figure 1). Now we address the results under ambient light. WT PA14 formed biofilms but the Δ*kinB* strain did not (Figure 4A). Our interpretation is that, in the WT, ambient light activates the BphP kinase and phosphotransfer to AlgB occurs. However, the opposing KinB phosphatase activity strips the phosphate from AlgB, thereby eliminating AlgB-dependent repression of biofilm formation. Thus, WT PA14 forms biofilms under ambient light. In the case of the Δ*kinB* mutant, since there is no KinB phosphatase present, ambient light is sufficient to drive BphP-mediated phosphorylation of AlgB, AlgB∼P accumulates, and it represses biofilm formation. Based on these results, we infer that the presence or absence of light can alter group behaviors such as biofilm formation in *P. aeruginosa*.

**Figure 4:**
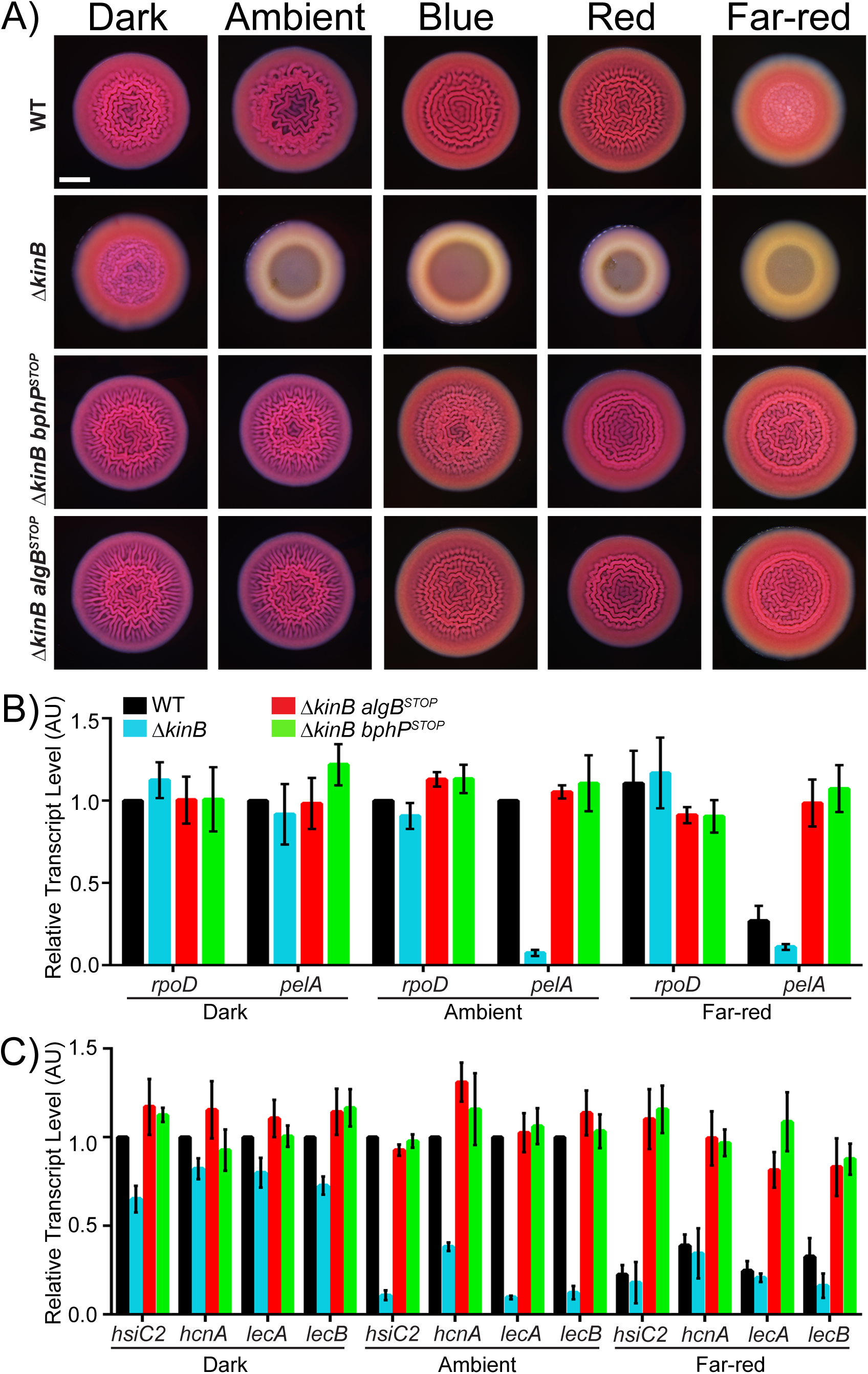
Photo sensing represses group behaviors in *P. aeruginosa*. A) Colony biofilm phenotypes are shown for WT PA14 and the designated mutants on Congo red agar medium after 72 h of growth under the indicated light conditions. Scale bar is 2 mm for all images. B) Relative expression of *rpoD* and *pelA* as measured by qRT-PCR in WT PA14 and the designated mutant strains grown as biofilms as in (A) in darkness, ambient light, and far-red light. C) Relative expression of *hsiC2*, *hcnA*, *lecA*, and *lecB* measured by qRT-PCR in WT PA14 and the designated mutants grown as biofilms as in (A) and light conditions as in (B). For panels B and C, data were normalized to 16S RNA levels and the WT levels were set to 1.0. AU denotes arbitrary units and error bars represent SEM for three biological replicates.

Ambient light is a composite of different wavelengths of light. The PA14 BphP bacteriophytochrome is reported to be a far-red light sensing HK *in vitro* (Tasler et al., 2005). We wondered if a particular wavelength of light could maximally activate the BphP kinase activity *in vivo*, and if so, perhaps, under that condition, the BphP kinase activity could override the KinB phosphatase, enabling light to repress biofilm formation in WT PA14. To test this notion, we exposed PA14 strains to blue, red, and far-red light and monitored biofilm formation. In contrast to WT PA14, the Δ*kinB* mutant failed to form biofilms under blue and red light, suggesting that BphP is a promiscuous photoreceptor that is activated by blue and red light (Figure 4A). Importantly, when WT PA14 was exposed to far-red light, it failed to make biofilms, but rather, exhibited the smooth phenotype identical to the Δ*kinB* mutant (Figure 4A). We conclude that far-red light is the preferred wavelength for BphP and is sufficient to repress biofilm formation in WT *P. aeruginosa*. Finally, we show that light-mediated repression of biofilm formation requires functional BphP and AlgB as both the Δ*kinB bphP^STOP^* and Δ*kinB algB^STOP^* mutants did not repress biofilm formation under the conditions tested (Figure 4A).

One mechanism by which light could suppress biofilm formation via BphP-AlgB is by down-regulating Pel production. To test this idea, we performed quantitative RT-PCR analyses on WT, Δ*kinB*, Δ*kinB algB^STOP^*, and Δ*kinB bphP^STOP^* biofilms in darkness and under ambient and far-red light and we quantified *pelA* transcript levels (Figure 4B). We used *rpoD* transcription as the control. Expression of *rpoD* did not change under any condition tested. Regarding *pelA*, analogous to what occurred for biofilm formation, there was no significant difference in *pelA* expression between the WT and the Δ*kinB* strain in the dark, whereas transcription of *pelA* was ∼14-fold lower in the Δ*kinB* strain than in the WT under ambient light. Repression of *pelA* expression depended on functional BphP and AlgB as the Δ*kinB bphP^STOP^* and Δ*kinB algB^STOP^* mutants transcribed *pelA* at high levels under both conditions. We conclude that dephosphorylation of AlgB does not occur in the Δ*kinB* mutant under ambient light. In this condition, BphP phosphorylates AlgB and AlgB∼P represses biofilm formation via down-regulation of *pelA* expression. Lastly, in the WT, *pelA* transcript levels were ∼4-fold lower under far-red light than in darkness. Therefore, far-red light is the strongest activator of BphP such that under far-red light, but not ambient light, the kinase activity of BphP overrides the phosphatase activity of KinB in the WT to drive AlgB∼P accumulation, repression of *pelA* expression, and consequently, repression of biofilm formation.

In Figure 2, we showed that BphP is required for AlgB-dependent repression of virulence gene expression. Our results in Figure 4 suggest that light, by controlling BphP-dependent phosphorylation of AlgB, could control virulence in *P. aeruginosa*. To explore this idea further, we quantified the expression of the virulence-associated genes, *hsiC2*, *hcnA*, *lecA*, and *lecB* in biofilms of WT PA14 and in the Δ*kinB*, Δ*kinB algB^STOP^*, and Δ*kinB bphP^STOP^* strains under darkness, ambient light, and far-red light. The results mirror those for biofilm formation and *pelA* transcription. Only in the absence of the opposing KinB phosphatase activity is ambient light sufficient to activate BphP, whereas far-red light-driven BphP kinase activity can override the KinB phosphatase activity allowing accumulation of AlgB∼P to levels that repress virulence gene expression. Again, light-mediated repression of virulence genes requires functional BphP and AlgB (Figure 4C). We conclude that BphP-dependent photo sensing represses virulence gene expression in *P. aeruginosa*.

Light possess both color (wavelength) and intensity properties. Above, we demonstrated that BphP can detect blue, red, and far-red light. To explore the possibility that *P. aeruginosa* BphP is also capable of detecting light intensity, we varied the intensity of far-red light since it has the most dramatic effect on PA14 phenotypes. We used repression of biofilm formation as the readout. Biofilm formation decreased with increasing intensity of far-red light in the WT and Δ*kinB* mutant but remained unaltered in the *bphP^STOP^* mutant (Figure 5A). The highest intensity of light we tested (bottom-most panel in Figure 5A) is similar to that present in natural sunlight (5.5 W/m^2^ in a 5 nm window around 730 nm; ASTM G173-03 Reference Solar Spectra, www.astm.org). At this intensity, WT biofilm formation was maximally repressed showing that BphP kinase dominates over KinB phosphatase. The Δ*kinB* mutant generates suppressor flares under this condition, suggesting that one role of the KinB phosphatase is to keep the BphP kinase activity in check. To verify that far-red light specifically altered biofilm behavior without affecting general physiology, we quantified *rpoD* and *pelA* transcript levels in the WT and *bphP^STOP^* mutant biofilm samples grown under the different light intensities (Figure 5B). Expression of *rpoD* did not change under any condition tested, while transcription of *pelA* decreased progressively in the WT with increasing intensity of far-red light. At the highest intensity of far-red light tested, expression of *pelA* was ∼12-fold lower than that in the *bphP^STOP^* mutant that cannot convey the light cue internally to AlgB. These results demonstrate that *P. aeruginosa* biofilm formation can be modulated simply by tuning the intensity of far-red light in which the strain is grown.

**Figure 5:**
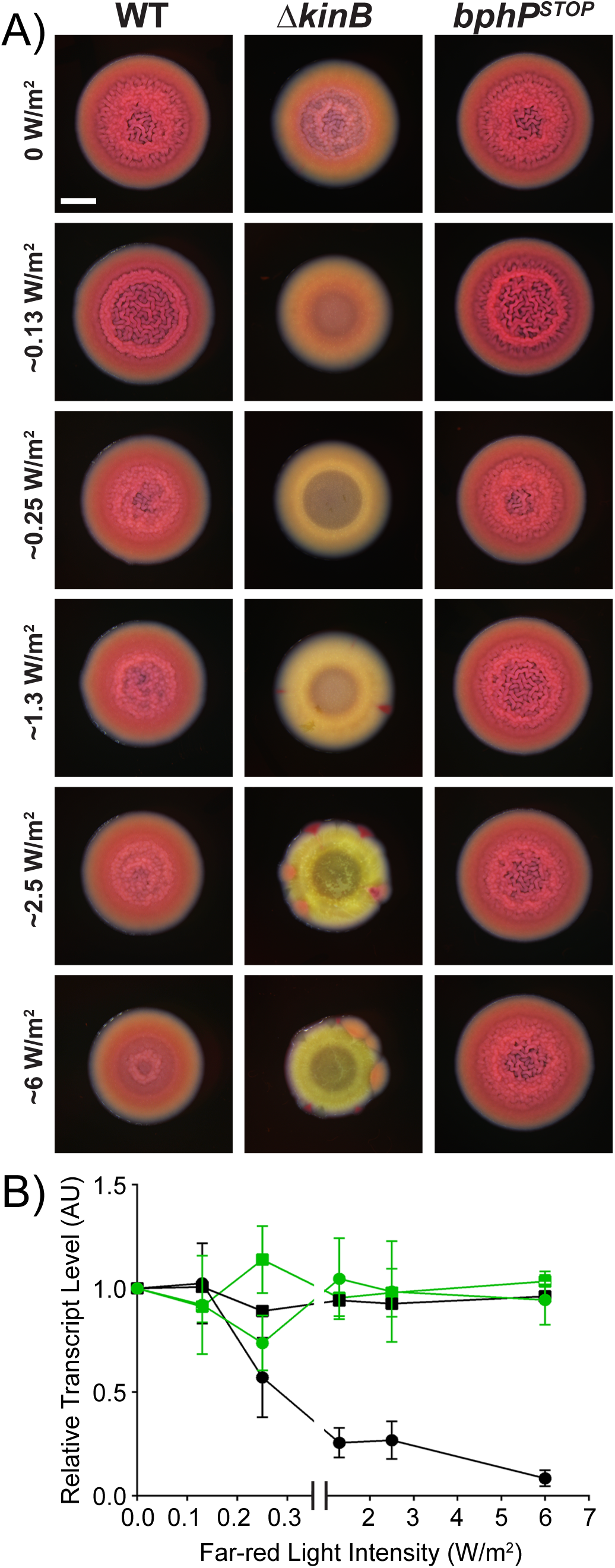
Far-red light intensity controls biofilm formation. A) Colony biofilm phenotypes are shown for WT PA14 and the designated mutants on Congo red agar medium after 72 h of growth under the indicated far-red light intensities. Scale bar is 2 mm for all images. B) Relative expression of *rpoD* (squares) and *pelA* (circles) measured by qRT-PCR in WT PA14 (black) and in the *bphP^STOP^* mutant (green) grown as biofilms as in (A). Data were normalized to 16S RNA levels and the WT levels at 0 mW/m^2^ far-red light were set to 1.0. AU denotes arbitrary units and error bars represent SEM for three biological replicates.

### The BphP-AlgB interaction is conserved in diverse bacteria

BphP bacteriophytochromes are a major class of photoreceptors widely distributed in non-photosynthetic bacteria (Gomelsky and Hoff, 2011). These BphP HKs either lack a partner RR, or when they are co-transcribed with a cognate RR gene, the physiological output of the circuit is unknown. Thus, the downstream signaling components and consequences of photo sensing in non-photosynthetic bacteria are not understood. Our discovery of AlgB as the cognate RR for the orphan light sensing BphP HK in a non-photosynthetic bacterium, coupled with our demonstration of the biofilm and virulence outputs of photo sensing, puts us in a position to test the generality of our findings. As a first step, we generated a phylogenetic tree containing 150 BphP orthologs that are the closest homologs to *P. aeruginosa* BphP (Figure 6A, S5). The majority of these BphP orthologs are present in non-photosynthetic bacteria from diverse phyla. The Pseudomonads fall into discrete clusters hinting at acquisition of BphP via horizontal gene transfer. With respect to AlgB and KinB, we find that, while KinB is present only in the Pseudomonads, *Acinetobacter baumannii*, and *Enterobacter cloacae* (Figure 6A, S1, S5), AlgB is present in ∼93% of the bacterial species in our BphP-based phylogenetic tree (Figure 6A, S5). We note that in all of the bacteria that do not encode AlgB, for example, *Deinococcus* spp., BphR is the cognate RR for BphP (Figure 6A, S1, 2, 5, and Bhoo et al., 2001). None of these *bphP*-encoding bacteria possesses both BphR and AlgB. Therefore, the pattern that emerges is that BphB is widely distributed in non-photosynthetic bacteria and the cognate RR is either AlgB or BphR.

**Figure 6:**
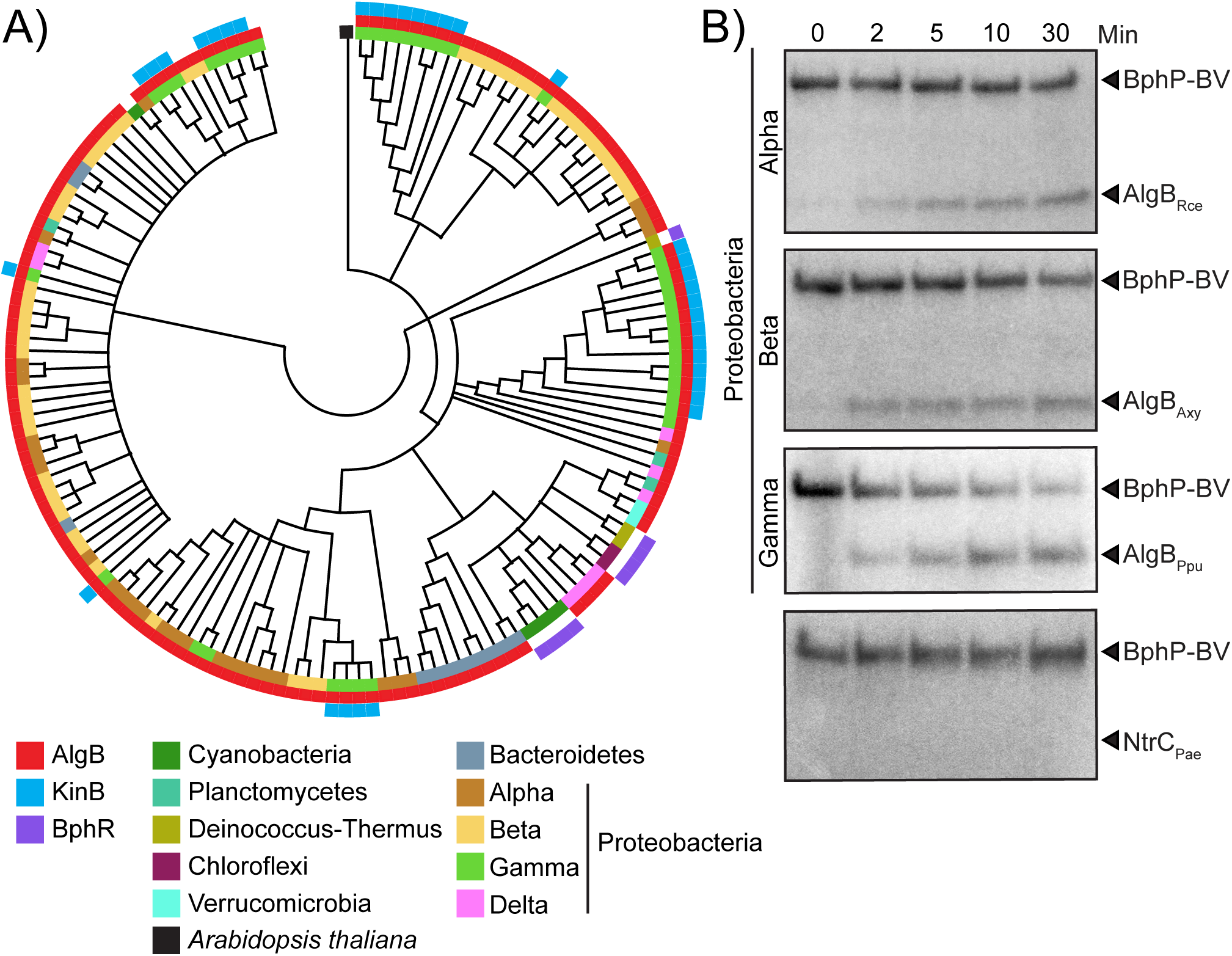
The BphP-AlgB interaction is conserved in diverse bacteria. A) Maximum likelihood-based phylogenetic tree for BphP showing the 150 closest orthologs to *P. aeruginosa* BphP, generated using MEGA-X software (Kumar et al., 2018). Co-occurrences of AlgB and KinB are depicted in red and blue, respectively. BphR is shown in purple. The other colors indicate bacterial phyla. The black square indicates *Arabidopsis thaliana* as the root of the tree. B) *In vitro* phosphorylation of AlgB orthologs from the α-Proteobacterium *Rhodospirillum centenum* (Rce), the β-Proteobacterium *Achromobacter xylosoxidans* (Axy), and the γ-Proteobacterium *Pseudomonas putida* (Ppu) by *P. aeruginosa* BphP-BV that had been autophosphorylated for 30 min. The bottom panel shows that phospho-transfer from *P. aeruginosa* P∼BphP-BV to *P. aeruginosa* NtrC does not occur.

To test if BphP can interact with and phosphorylate AlgB in bacteria other than *P. aeruginosa*, we purified AlgB orthologs from diverse Proteobacteria: *Rhodospirillum centenum* (α), *Achromobacter xylosoxidans* (β) and *Pseudomonas putida* (γ). We incubated these AlgB proteins with an equimolar concentration of autophosphorylated *P. aeruginosa* BphP-BV. Phosphotransfer from BphP-BV to the AlgB orthologs occurred in all cases, albeit to varying degrees (Figure 6B). To eliminate the possibility that BphP-BV is a promiscuous kinase for NtrC family RRs, we purified NtrC from *P. aeruginosa* and incubated with autophosphorylated BphP-BV. BphP-BV failed to phosphorylate NtrC (Figure 6B). We conclude that BphP is the specific HK for AlgB, and AlgB appears to have a conserved function in photosensory signal transduction in diverse bacteria.

## DISCUSSION

Our study reveals that the non-photosynthetic pathogenic bacterium *P. aeruginosa* detects and responds to light to repress group behaviors including virulence factor production and biofilm formation. The photoreceptor BphP functions as a light-activated HK that phosphorylates the AlgB RR. AlgB∼P represses group behaviors but is antagonized by its canonical HK KinB. Specifically, KinB dephosphorylates AlgB, and thus, KinB functions as an activator of group behaviors. Our work shows that AlgB functions as a hub protein that has three inputs -- quorum sensing via RhlR, photo sensing via BphP, and an unknown signal via KinB. While quorum sensing activates *algB* expression, photo sensing activates AlgB function, and thus the presence or absence of light can override the quorum-sensing input from RhlR. We reason that, at high cell density, RhlR will drive AlgB production. However, if there is no light, BphP will not phosphorylate and activate AlgB. In turn, AlgB will not repress group behaviors. To our knowledge, the BphP-AlgB photosensory signal transduction cascade represents the first example of light-mediated control of group behaviors in the global pathogen *P. aeruginosa*.

Light is a ubiquitous source of energy that drives the anabolic process of photosynthesis in photosynthetic organisms. However, the wide distribution of photoreceptors in all domains of life suggests roles for photo sensing in behaviors far beyond photosynthesis. Plants, for example, use light cues to regulate activities such as seed germination (Mathews, 2006), stomatal opening (Shimazaki et al., 2007), and defenses against microbes (Bhardwaj et al., 2011; Neukermans et al., 2012; Roden and Ingle, 2009). Furthermore, plant vascular systems can function as bundles of optical fibers to efficiently transmit light, particularly far-red light, that is not absorbed by plant pigments, allowing opportunities for photo sensing in roots and possibly in the rhizosphere (Lee et al., 2016). Many of the *bphP*-encoding bacteria from the phylogenetic tree in Figure 6A that also possess AlgB, are members of the rhizosphere microbiome (Duran et al., 2018). Perhaps these non-photosynthetic bacteria exploit light cues to colonize and/or to fine-tune their mutualistic or pathogenic interactions with their plant hosts as well as adjust their physiology in the rhizosphere.

Light provides spatial and temporal information to higher organisms. Does light serve a similar purpose in bacteria? Recent studies have reported that BphP plays a role in multiple stages of infection by the foliar plant pathogens *Xanthomonas campestris pv. campestris* and *Pseudomonas syringae pv. syringae* (Bonomi et al., 2016; Wu et al., 2013), in each case, via an unknown but putative downstream RR. Based on our phylogenetic analysis, we speculate that AlgB fulfils this role. We further speculate that *P. aeruginosa*, which is a plant pathogen (Starkey and Rahme, 2009), responds to light cues via the BphP-AlgB TCS to appropriately modulate its biofilm and virulence programs, particularly, to inhibit virulence during daylight enabling avoidance of plant defense mechanisms. For instance, during the day, chlorophyll in leaves removes most of the red wavelength from sunlight but little of the far-red spectrum (Smith, 2000). Thus, far-red light is readily available, and based on our work here, could signal to *P. aeruginosa* to tamp down virulence factor production and biofilm formation, allowing it to optimize those programs in line with host conditions as shaded leaves are more susceptible to infection than leaves exposed to direct light (de Wit, 2013).

In addition to providing spatial-temporal information, light can also reveal other key parameters to which bacteria respond. Photoreceptors fall into six families depending on the structure of the light-absorbing chromophore: rhodopsins, xanthopsins, cryptochromes, LOV domain-containing phototropins, blue-light sensing using flavin (BLUF)-domain proteins, and phytochromes (Kottke et al., 2018; Shcherbakova et al., 2015). Detection of blue light via LOV and BLUF domain proteins modulates general stress responses in some non-photosynthetic bacteria such as *Bacillus subtilis* and *Caulobacter crescentus* (Ávila-Pérez, et al., 2006; Purcell et al., 2007). Light, through the LOV-HK of the mammalian pathogen *Brucella abortus* is crucial for virulence in a macrophage infection model, although the components connecting light to the virulence response remain undefined. It is also proposed that *B. abortus* uses light as an indicator of whether it is inside or outside of its animal host (Swartz et al., 2007). The *P. aeruginosa* genome does not encode LOV or BLUF domain proteins (Horst et al., 2007). *P. aeruginosa*, possesses only one identifiable photoreceptor, BphP (Horst et al., 2007). Nonetheless, we showed that *P. aeruginosa* is capable of detecting blue, red, and far-red light via BphP (Figure 4A). Perhaps, an advantage of BphP promiscuity is that it enables detection of higher energy, and therefore, phototoxic blue light, in addition to the lower energy but highly penetrative far-red light. Such a scenario would endow *P. aeruginosa* with the plasticity to diversify its physiological outputs in response to particular wavelengths of light, without the necessity of a distinct photoreceptor for each wavelength. We do not yet know the molecular mechanisms that permit BphP to detect blue, red, and far-red light, whether there are one or multiple chromophores involved, and whether there exist different output regulons for different input light wavelengths.

An advantage *P. aeruginosa* could accrue by sensing light on or within a mammalian host would be the ability to tune into the host circadian rhythm and its associated responses. Circadian clocks influence various aspects of health and disease such as sleep/wake cycles and metabolism (Curtis et al., 2014; Scheiermann et al., 2013). Disruption of circadian rhythms are associated with fitness costs (Scheiermann et al., 2013). In mammals, both innate and adaptive immune responses are controlled by the circadian clock such that the immune system is primed to combat pathogens during the host active phase while immune functions undergo regeneration and repair during the resting phase of the daily cycle. Parasites such as *Plasmodium* spp. that cause malaria, synchronize their replication cycle with host circadian rhythms for optimized infection and dissemination (Donnell et al., 2011). Likewise, viruses such as Herpes and Influenza A have been shown to exploit the mammalian circadian clock for their own gain i.e., to successfully avoid host immune responses enabling maximal replication (Edgar et al., 2016; Sundar et al., 2015). Furthermore, the human body emits light, albeit at 1,000-fold lower intensity than is visible to the naked eye, but intriguingly, photon emission peaks during the day and is lowest at night, and therefore, might be controlled by the endogenous circadian clock (Kobayashi et al., 2009). Perhaps, *P. aeruginosa* uses light as a signal that reveals when the host immune response is at peak function, and accordingly, at that time, *P. aeruginosa* represses biofilm formation and virulence factor expression as a mechanism that enhances evasion of host defenses. If so, a human host infected with *P. aeruginosa* during the night would be colonized to higher levels, compared to a host acquiring an infection during the day. Synchronizing infectivity with light/dark cues to enable optimal infection could be a common feature of non-photosynthetic photoreceptor-harboring pathogens.

*P. aeruginosa* is a priority pathogen on the CDC (Centers for Disease Control and Prevention) ESKAPE pathogen list (a set of bacteria including *Enterococcus faecium, Staphylococcus aureus, Klebsiella pneumoniae, Acinetobacter baumannii, P. aeruginosa* and *Enterobacter* spp. that are designated as multi-drug resistant pathogens requiring new antimicrobials for treatment), and a critical pathogen on the WHO (World Health Organization) priority list (Rice, 2008; Pendleton et al., 2013; Tacconelli et al., 2017). Our phylogenetic analysis suggests that the KinB-AlgB-BphP module is conserved in the genomes of *A. baumannii* and *Enterobacter cloacae*, perhaps acquired from *P. aeruginosa* via horizontal gene transfer, as the AlgB primary sequence is nearly identical between the three species. We speculate that, beyond *P. aeruginosa*, BphP-AlgB-dependent photo sensing also affects the physiology, and possibly the virulence of these ESKAPE pathogens. Collectively, the results from this study provide unanticipated insight into *P. aeruginosa* physiology and a surprising possibility for therapeutic intervention—shining light on a deadly and actively studied pathogen, *P. aeruginosa*, to attenuate virulence and biofilm formation.

## Supporting information

Mukherjee Supplemental Information

## ACKNOWLEDGEMENTS

We thank Wei Wang and the Genomics Core Facility at Princeton University for help with whole genome sequencing. We thank Ned Wingreen, Anne-Florence Bitbol, Joseph E. Sanfillipo, and all members of the Bassler group for thoughtful discussions. This work was supported by the Howard Hughes Medical Institute, NIH Grant 5R37GM065859, and National Science foundation Grant MCB-1713731 to B.L.B., and a NIH Grant 1K99GM129424-01 to S.M.

## AUTHOR CONTRIBUTIONS

S.M., V.S. and M.J. conducted experiments; S.M. and M.T. analyzed data; S.M., M.J. and B.L.B. designed the experiments; S.M. and B.L.B. wrote the paper.

## MATERIALS AND METHODS

### Bacterial strains and growth conditions

All strains and plasmids used in this study are listed in Supplemental Tables S3 and S4, respectively. *P. aeruginosa* PA14 and mutants were grown at 37°C in lysogeny broth (LB) (10 g tryptone, 5 g yeast extract, 5 g NaCl per L), in 1% Tryptone broth (TB) (10 g tryptone per L), or on LB plates fortified with 1.5% Bacto agar. When appropriate, antimicrobials were included at the following concentrations: 400 µg/mL carbenicillin, 30 µg/mL gentamycin, and 100 μg/mL irgasan. *Escherichia coli* was grown at 37°C in LB, or on LB plates fortified with 1.5% Bacto agar and the following concentrations of antimicrobials as appropriate: 15 µg/mL gentamycin, 50 µg/mL kanamycin, and 100 μg/mL ampicillin. Isopropyl β-D-thiogalactopyranoside (IPTG, Sigma) was added to the medium at the indicated concentrations when appropriate.

### Mutant strain and plasmid construction

Strains and plasmids were constructed as described previously (Mukherjee et al. 2017). Briefly, to construct marker-less in-frame chromosomal deletions and substitutions in PA14, DNA fragments flanking the gene of interest were amplified, assembled by the Gibson method (Gibson et al., 2009), and cloned into suicide vector pEXG2 (Hmelo et al., 2015). The resulting plasmids were used to transform *E. coli* SM10λ*pir*, and subsequently, mobilized into PA14 strains via biparental mating. Exconjugants were selected on LB containing gentamicin and irgasan, followed by recovery of deletion mutants on LB medium containing 5% sucrose. Candidate mutants were confirmed by PCR and Sanger sequencing. Transposon insertions in the PA14 chromosome were generated by mating the PA14 Δ*rhlR* parent strain with *E. coli* SM10λ*pir* harboring pIT2 (IS*lacZ*/*hah*) (Jacobs et al., 2003). Insertion mutants were selected on LB agar containing 60 µg/mL tetracycline and 100 μg/mL irgasan was included in the agar to counter select against the *E. coli* donor. Transposon insertion locations were determined by arbitrary PCR and sequencing as described previously (Jacobs et al., 2003).

Protein production constructs were generated by amplifying the *algB*, *kinB*, and *bphP* coding regions and cloning them in pET28b or pET21b expression vectors (Novagen) to obtain pET28b-His6-AlgB, pET21b-KinB-His6, and pET21b-BphP-His6, respectively. To generate the AlgB^D59N^, KinB^P390S^, and BphP^H513A^ variants, the corresponding mutations were engineered on to the pET28b-His6-AlgB, pET21b-KinB-His6, and pET21b-BphP-His6 plasmids, respectively, via Gibson assembly. AlgB orthologs from *R. centenum* (Rce) and *A. xylosoxidans* (Axy) were amplified from gene fragments obtained from Integrated DNA Technologies, and that from *P. putida* (Ppu) was amplified from the *P. putida* KT2440 genome. All of the gene orthologs were cloned into the pET28b plasmid.

### Pyocyanin assay

PA14 strains were grown overnight in LB liquid medium at 37°C with shaking at 250 rotations per minute (rpm). The cells were pelleted by centrifugation at 21,130 × g for 2 min and the clarified supernatants were passed through 0.22 μm filters (Millipore) into clear plastic cuvettes. The OD_695_ of each sample was measured on a spectrophotometer (Beckman Coulter DV 730) and normalized to the culture cell density which was determined by OD_600_.

### Colony biofilm assay

The procedure for establishing colony biofilms has been described (Mukherjee et al., 2017). Briefly, 1 μL of overnight cultures of PA14 strains were spotted onto 60 × 15 mm Petri plates containing 10 mL 1% TB medium fortified with 40 mg/L Congo red and 20 mg/L Coomassie brilliant blue dyes and solidified with 1% agar. Biofilms were grown at 25°C for 72 h in an incubator (Benchmark Scientific) and images were acquired using a Leica stereomicroscope M125 mounted with a Leica MC170 HD camera at 7.78x zoom.

For biofilms exposed to specific wavelengths of light, the following light-emitting diodes (LED) were used: blue – 430 nm (Diffused RGB LED, #159, Adafruit), red – 630 nm (Diffused RGB LED, #159, Adafruit), and far-red – 730 nm (LST1-01G01-FRD1-00, Opulent). Ambient light exposure refers to biofilms grown under laboratory light conditions. For the biofilms shown in Figure 4A, light intensity was normalized by photon flux and the following intensities were used: blue (0.7 W/m^2^), red (1 W/m^2^), and far-red (1.1 W/m^2^). Light intensity was calibrated using a laser power meter (Ophir) in a 5 nm window at the appropriate wavelength. Biofilm samples were grown in custom laser-cut acrylic chambers. Each chamber housed a single LED light source and an individual petri plate containing 4 technical replicates. Samples exposed to darkness were housed in the same chambers as the light-exposed samples, but with no current applied to the LEDs.

### qRT-PCR

WT PA14 and mutant strains were harvested from planktonic cultures (OD_600_ = 1.0) or from biofilms grown for 72 h. RNA was purified using the Zymo Research kit, and the preparations were subsequently treated with DNAse (TURBO DNA-free™, Thermo Fisher). cDNA was synthesized using SuperScript® III Reverse Transcriptase (Invitrogen) and quantified using PerfeCTa® SYBR® Green FastMix® Low ROX (Quanta Biociences).

### Protein purification. His6-AlgB

The pET28b-His6-AlgB protein production vector was transformed into *E. coli* BL21 (DE3) and the culture grown to ∼0.8 OD_600_ in 1 L of LB supplemented with 50 μg/mL kanamycin at 37°C with shaking at 220 rpm. Protein production was induced by the addition of 1 mM IPTG, followed by incubation of the culture for another 3 h at 25°C with shaking. The cells were pelleted by centrifugation at 16,100 × g for 20 min and resuspended in AlgB-lysis buffer [50 mM NaH_2_PO_4_ pH 8.0, 300 mM NaCl, 1 mM MgCl_2_, 1 mM DTT, 5% glycerol, 0.1% Triton X-100, 10 mM Imidazole, and protease inhibitor cocktail (Roche)]. The preparation was frozen at −80°C overnight. The frozen cell pellet was thawed on ice and the cells lysed by sonication (1 s pulses for 15 s). The sample was subjected to centrifugation at 32,000 × g for 30 min at 4°C. The resulting clarified supernatant was combined with Ni-NTA resin (Novagen) and incubated for 3 h at 4°C. The bead/lysate mixture was loaded onto a 1 cm separation column (Bio-Rad), the resin was allowed to pack, and then it was washed with AlgB-wash buffer [50 mM NaH_2_PO_4_ pH 8.0, 300 mM NaCl, 1 mM MgCl_2_, 1 mM DTT, 5% glycerol, 0.1% Triton X-100, 30 mM Imidazole, and protease inhibitor cocktail (Roche)]. Resin-bound His6-AlgB was eluted twice with 1 mL AlgB-wash buffer containing 250 mM Imidazole. Fractions were analyzed by SDS-PAGE and the gel was stained with Coomassie Brilliant Blue to assess His6-AlgB purity. Purified protein was dialyzed in AlgB-storage buffer [50 mM NaH_2_PO_4_ pH 8.0, 300 mM NaCl, 1 mM MgCl_2_, 1 mM DTT, 5% glycerol, and 0.1% Triton X-100], and stored at −80°C.

### BphP-His6

The pET21b-BphP-His6 protein production vector was transformed into *E. coli* BL21-CodonPlus (DE3)-RIPL (Agilent Technologies). BphP-His6 was purified as described for His6-AlgB with the following changes in buffers: BphP-lysis buffer [50 mM NaH_2_PO_4_ pH 8.0, 300 mM NaCl, 1% Triton X-100, 0.1% β-mercaptoethanol, 10 mM Imidazole, and protease inhibitor cocktail (Roche)], BphP-wash buffer [50 mM NaH_2_PO_4_ pH 8.0, 300 mM NaCl, 1% Triton X-100, 0.1% β-mercaptoethanol, 30 mM Imidazole, and protease inhibitor cocktail (Roche)], and BphP-storage buffer [50 mM NaH_2_PO_4_ pH 8.0, 300 mM NaCl, 1% Triton X-100, 0.1% β-mercaptoethanol, 5% glycerol].

### KinB-His6

The pET21b-KinB-His6 protein production vector was transformed into *E. coli* BL21 (DE3). KinB-His6 protein was purified exactly as described above for BphP-His6.

### Phosphorylation assays

Autophosphorylation assays were performed with purified WT BphP and the BphP^H513A^ variant or with KinB and the KinB^P390S^ variant. 100 µM BphP or BphP^H513A^ was incubated under ambient light with 10-fold molar excess of Biliverdin (Sigma-Aldrich) for 1 h prior to the assay to form the light-activated BphP-BV stocks. Reactions were carried out in phosphorylation buffer [50 mM Tris pH 8.0, 100 mM KCl, 5 mM MgCl_2_, and 10% (v/v) glycerol], and were initiated with the addition of 100 µM ATP and 2 µCi [γ-^32^P]-ATP (Perkin Elmer). Reactions were incubated at room temperature and terminated by the addition of SDS-PAGE loading buffer. Reaction products were separated using SDS-PAGE. Gels were dried at 80°C on filter paper under vacuum, exposed to a phosphoscreen overnight, and subsequently analyzed using a Typhoon 9400 scanner and ImageQuant software. For phosphotransfer to AlgB, an equimolar concentration of AlgB was added to the phospho-BphP-BV or phospho-KinB proteins. Reactions were incubated at room temperature for the indicated times and terminated by the addition of SDS-PAGE loading buffer.

Dephosphorylation of AlgB∼P: 10 µM AlgB was phosphorylated for 30 min in reactions containing 10 µM BphP-BV, 100 µM ATP, and 2 µCi [γ-^32^P]-ATP in phosphorylation buffer. Subsequently, the reactions containing AlgB∼P were applied to gel filtration spin columns (Probe Quant G-50, GE healthcare) to remove ATP. Dephosphorylation reactions were initiated by adding 10 µM KinB or KinB^P390S^. Aliquots were taken at the indicated times and analyzed as described above.

### Phos-Tag SDS-PAGE and Western Blotting

WT PA14 and mutant strains were harvested from planktonic cultures (OD_600_ = 1.0). Cells were resuspended in 100 μl of ice-cold BugBuster reagent (Novagen) containing EDTA-free Protease Inhibitor Cocktail (Roche), followed by end-over-end rotation on a nutator at room temperature for 30 min. Cell debris was removed by centrifugation (4°C at 10,000 rpm for 1 min). 50 μL of 4x SDS-PAGE loading buffer (Thermo Fisher Scientific) containing 15% β-mercaptoethanol was combined with 50 μL of the sample supernatant. 10 μL of samples were loaded onto a 12.5% SuperSepTM Phos-tag^TM^ gel (Wako Pure Chemical Industries). Samples were subjected to electrophoresis at 4°C for 3 h. Gels were incubated for 20 min on a shaking platform in 1x transfer buffer containing 1 mM EDTA, and re-equilibrated for 20 min in 1x transfer buffer lacking EDTA. Proteins were transferred to nitrocellulose membranes, blocked with 5% skim milk in TBS at room temperature for 1 h, and incubated with primary anti-FLAG antibody (Sigma Aldrich) at 1:5000 dilution in 5% skim milk in TBS overnight 4°C on a rocking platform. Membranes were washed three times with TBS-Tween 20 at room temperature for 10 min, on a rocking platform, and subsequently developed with SuperSignal West Femto Kit (Thermo Scientific) and captured with LAS-4000 Imager (GE Healthcare).

### Whole genome sequencing

*P. aeruginosa* strains were harvested from planktonic cultures (OD_600_ = 2.0) and DNA was purified using DNeasy Blood & Tissue kit (Qiagen). The Nextera DNA Library Prep kit (Illumina, CA) was employed with 2 ng of genomic DNA to prepare the library. Unique barcodes were added to each sample to enable multiplexing. The libraries were examined for quality using Bioanalyzer DNA High Sensitivity chips (Agilent, CA) and quantified using a Qubit fluorometer (Invitrogen, CA). DNA libraries from the different strains were pooled at equal molar amounts and sequenced using an Illumina MiSeq as pair-end 2×100 nt reads. Only the Pass-Filter (PF) reads were used for further analysis.

Whole-genome sequencing data were processed with breseq version 0.33.2 to identify mutations relative to the reference *P. aeruginosa* UCBPP-PA14 genome (www.pseudomonas.com; Winsor et al., 2016). All high-confidence and putative SNPs and deletion events were confirmed by a manual examination of the read pileups with GenomeViewer IGV 2.4.8. A sample collected prior to the suppressor mutation screen was aligned against the reference genome of PA14, yielding a manually curated list of 25 differences acquired by our laboratory strain prior to the experiment (19 SNPs, 6 single-nucleotide indels). Applying these differences to PA14 using gdtools (part of the breseq package) yielded an updated reference genome against which all other samples were compared. Table S2 reports all high-confidence mutations identified in this analysis.

