## Supplementary material for "Photo sensing and quorum sensing are integrated to control bacterial group behaviors": Mukherjee Supplemental Information

#### SUPPLEMENTAL FIGURE LEGENDS AND TABLE TITLES

**Supplemental Figure 1: Gene conservation for KinB, AlgB, BphO, and BphP.** The genes flanking *kinB, algB, bphO*, and *bphP* are diagrammed for the indicated genomes. The relative gene positions and orientations are accurate, but gene lengths are not to scale.

**Supplemental Figure 2: Multiple sequence alignment for AlgB orthologs.** Primary sequence alignment of NtrC (first line) and AlgB (second line) from *P. aeruginosa* (Pae), and AlgB orthologs (third through twelfth lines) from *Pseudomonas fluorescens* (Pfl), *Pseudomonas syringae* (Psy), *Pseudomonas protegens* (Ppr), *Pseudomonas stutzeri* (Pst), *Pseudomonas entomophila* (Pen), *Pseudomonas putida* (Ppu), *Acinetobacter baumannii* (Aba), *Enterobacter cloacae* (Ecl), *Achromobacter xylosoxidans* (Axy), *Rhodospirillum centenum* (Rce), and BphR (thirteenth line) from *Deinococcus radiodurans.* Highly conserved amino acids are highlighted in black. Residue 59 is shown by the green asterisk. The GAFTA motif required for interaction with $\sigma^{54}$ is indicated by the magenta line.

**Supplemental Figure 3: AlgB$^{D59N}$, KinB$^{P390S}$ and BphP$^{H513A}$ are produced and stable in *P. aeruginosa*.** A) Western blot analysis of whole cell lysates from the indicated strains, all of which have the *algB$^{STOP}$* allele at the native locus in the genome and carry an empty vector or *3xFLAG-algB or 3xFLAG-algB$^{D59N}$* on the pBBR1-MCS5 plasmid under the $P_{lac}$ promoter. The same cell lysates were probed for RNAP as the loading control. B) Colony biofilm phenotypes of WT PA14, and the Δ*kinB*, Δ*kinB algB$^{STOP}$*, and Δ*kinB 3xFLAG-algB* mutants. Scale bar is 2 mm. C) SDS-PAGE analysis of whole cell lysates from the indicated strains. The gel was stained for SNAP using SNAP-Cell® 647-SiR fluorescent substrate (NEB). Lysozyme was added as the loading control. D) Colony biofilm phenotypes of the WT, *kinB-SNAP*, and *kinB$^{P390S}$-SNAP* strains. Scale bar is 2 mm. E) Western blot analysis of whole cell lysates from the indicated strains. The same cell lysates were probed for RNAP as the loading control.

**Supplemental Figure 4: BphP$^{H513A}$-BV cannot but KinB and KinB$^{P390S}$ can phosphorylate AlgB *in vitro*.** A) Autophosphorylation of BphP-BV was carried out for 30 min (left most lane) followed by addition of AlgB (second lane) or AlgB$^{D59N}$ (third lane) for 30 min. The kinase-defective BphP$^{H513A}$-BV was incubated with radiolabeled ATP for 30 min (fourth lane), followed by addition of AlgB (fifth lane) B) Autophosphorylation of KinB was carried out for 30 min and samples were removed at the indicated times. C) An equimolar amount of AlgB was added to KinB that had been autophosphorylated for 30 min as in (B). Samples were taken at the indicated times. D and E) As in A and B, respectively, but for the phosphatase-deficient protein KinB$^{P390S}$.

**Supplemental Figure 5: The BphP-AlgB interaction is conserved in diverse bacteria.** Enlarged maximum likelihood-based phylogenetic tree for BphP from Figure 6A showing the 150 closest orthologs to *P. aeruginosa* BphP. Co-occurrences of AlgB and KinB are depicted using red and blue dots, respectively. The presence of BphR is shown by purple dots. The colored squares indicate the corresponding bacterial phyla. The black square indicates *Arabidopsis thaliana* as the root of the tree.

**TABLE S1: Transposon insertion locations**

**TABLE S2: Suppressor mutations of the Δ*kinB* smooth biofilm phenotype**

**TABLE S3: Bacterial strains**

**TABLE S4: Plasmids**

### Supplemental Figure 1

A) *algB-kinB* genetic neighborhood — *bphP* genetic neighborhood

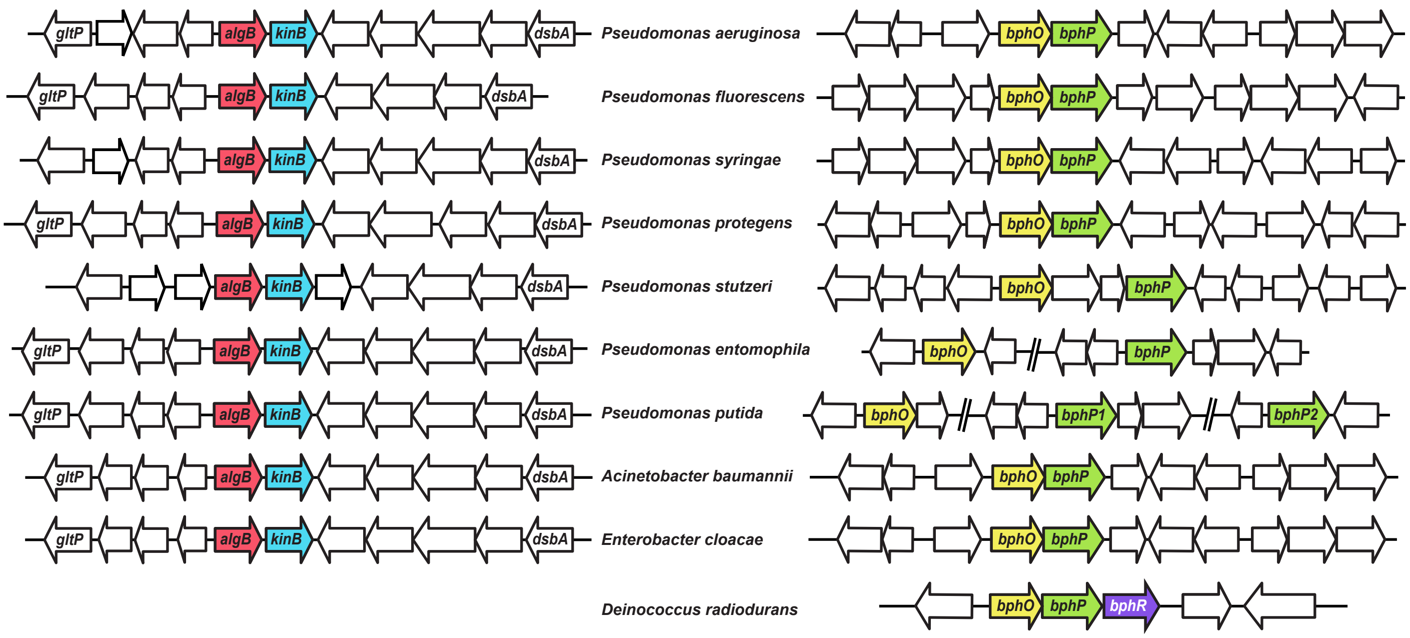

### Supplemental Figure 2

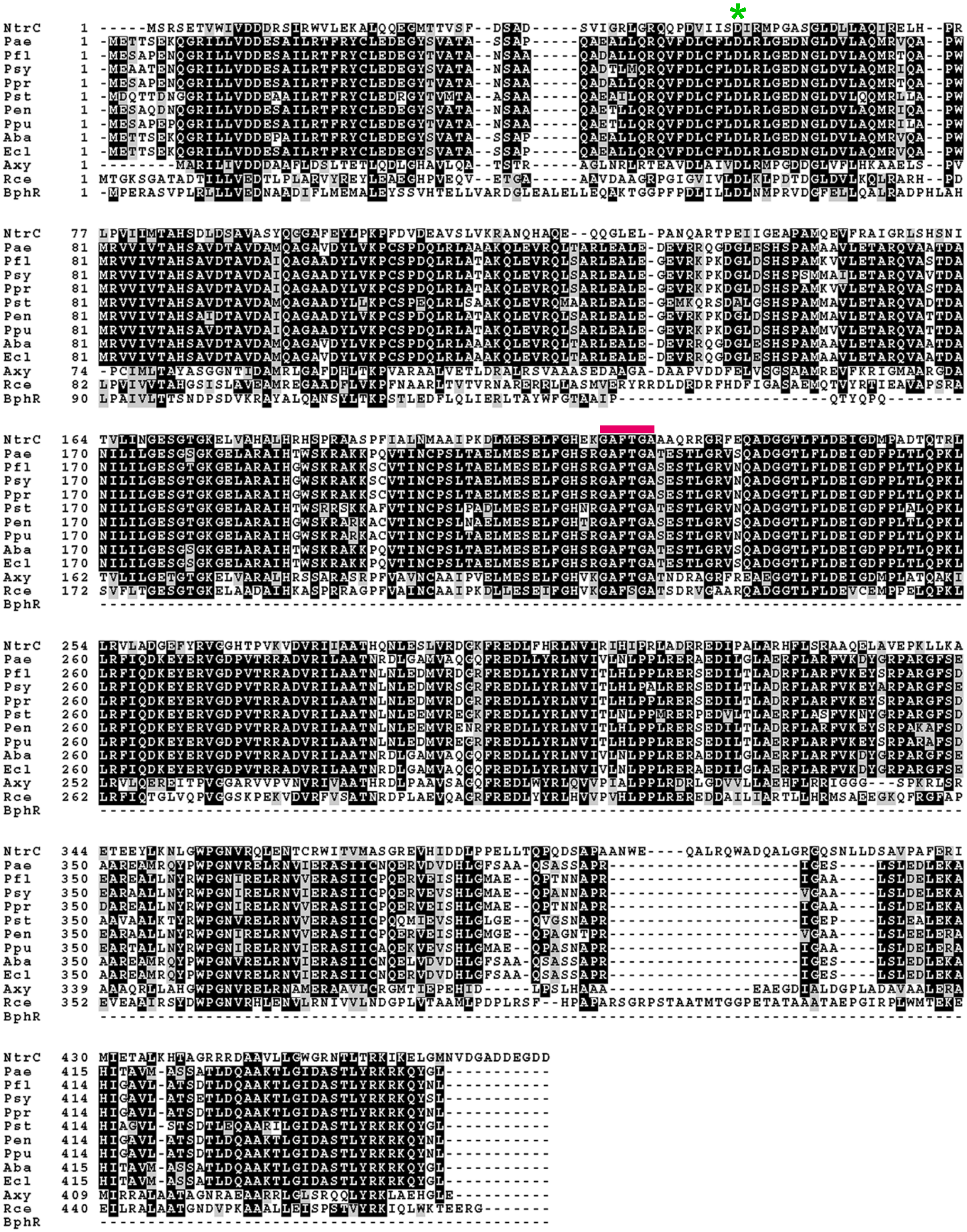

### Supplemental Figure 3

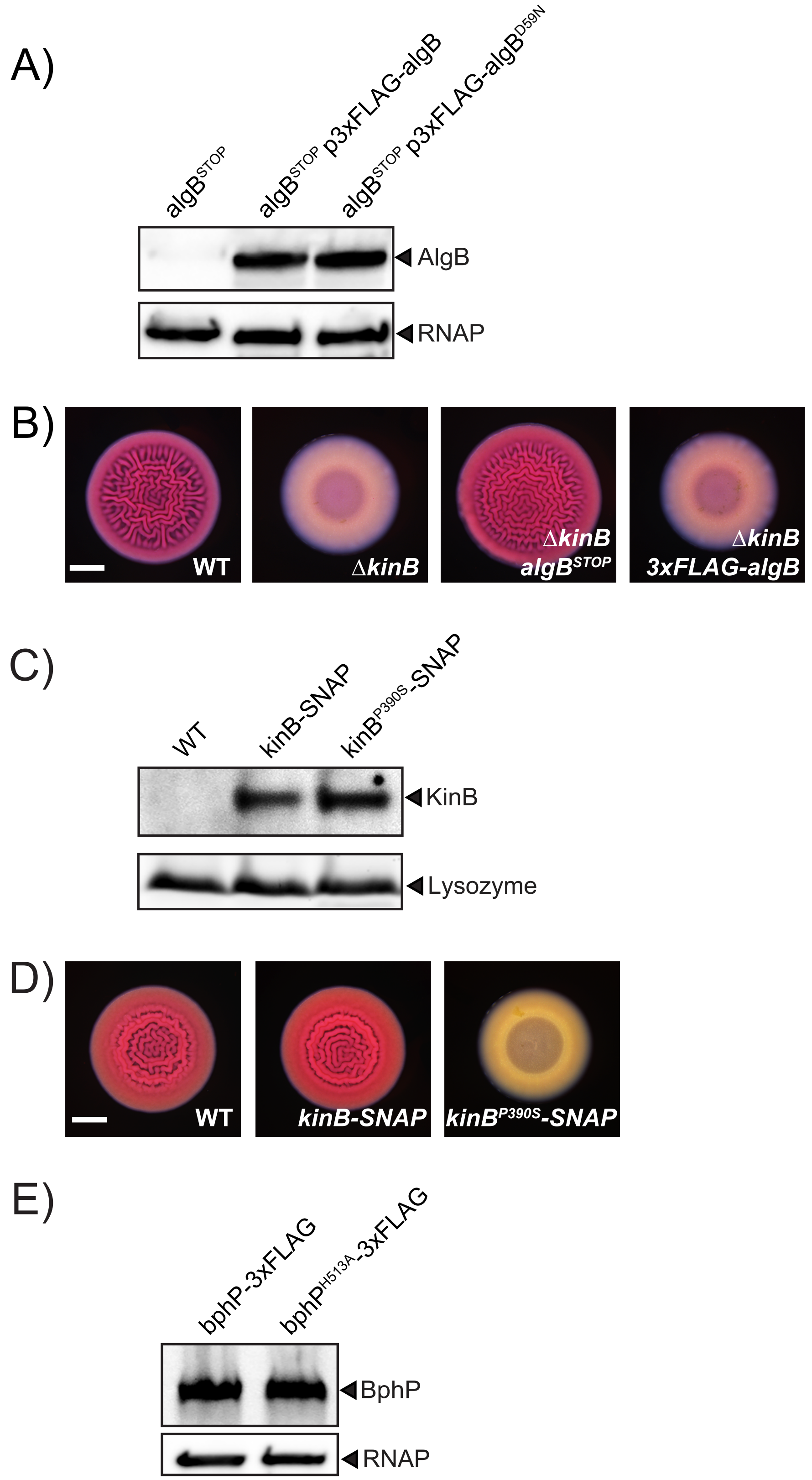

### Supplemental Figure 4

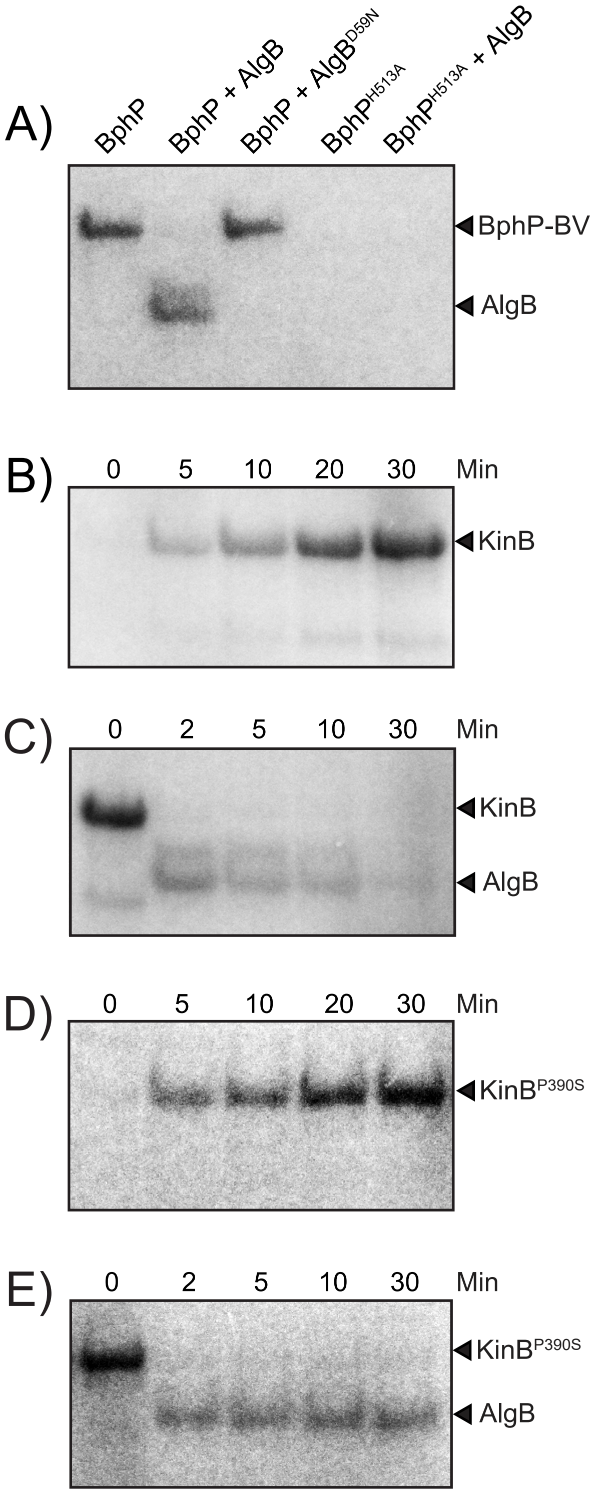

### Supplemental Figure 5

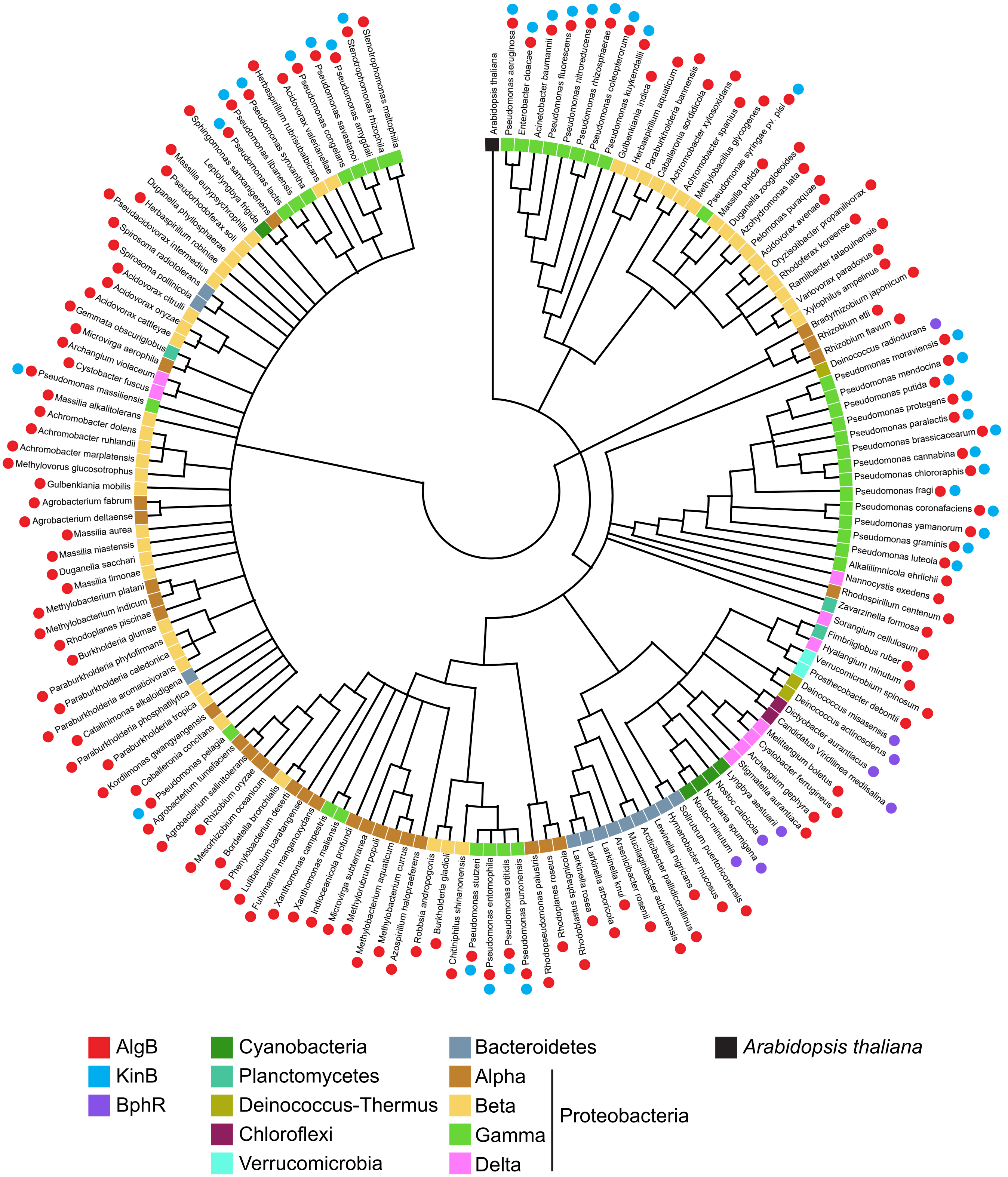

**TABLE S1: Transposon insertion locations**

| PA14 ID[a] | Gene name and description of encoded protien |
|---|---|
| PA14_19090 | *dcd*, deoxycytidine triphosphate deaminase |
| PA14_22060 | hypothetical protein |
| PA14_23420 | *zbdP,* zinc-binding dehydrogenase |
| PA14_23470 | *wbpM*, nucleotide sugar epimerase/dehydratase |
| PA14_24480 | *pelA*, extracellular polysaccharide biosynthesis protein |
| PA14_24490 | *pelB*, extracellular polysaccharide biosynthesis protein |
| PA14_32610 | *dsbG*, disulfide isomerase/thiol-disulfide oxidase |
| PA14_33270 | *pvdG*, pyoverdine synthetase |
| PA14_33700 | *pvdF*, pyoverdine synthetase |
| PA14_35720 | hypothetical protein |
| PA14_38510 | *hmgA*, homogentisate 1,2-dioxygenase |
| PA14_40860 | hypothetical protein |
| PA14_44070 | *gltA*, type II citrate synthase |
| PA14_52060 | hypothetical protein |
| PA14_58760 | *pilC*, type 4 fimbrial biogenesis protein |
| PA14_59630 | hypothetical protein |
| PA14_59800 | *pvrS*, two-component sensor kinase |
| PA14_66620 | *pilQ*, type 4 fimbrial biogenesis protein |
| PA14_66660 | *pilM*, type 4 fimbrial biogenesis protein |
| PA14_72390 | *kinB*, two-component sensor kinase |

a: annotation from www.pseudomonas.com (Winsor et al., 2016)

**TABLE S2: Suppressor mutations of the Δ*kinB* smooth biofilm phenotype**

| Suppressor | PA14 ID[a] | Gene name | Nucleotide position | Mutation |
|---|---|---|---|---|
| SM1045 | PA14_72380 | *algB* | 6447033 | Δ10 bp |
| SM1062 | PA14_72380 | *algB* | 6447033 | Δ10 bp |
| SM1063 | PA14_72380 | *algB* | 6447033 | Δ10 bp |
| SM1064 | PA14_72380 | *algB* | 6447033 | Δ10 bp |
| SM1067 | PA14_10700 | *bphP* | 919652 | Δ1603 bp |
| SM1068 | PA14_72380 | *algB* | 6447064 | Δ1 bp |
| SM1072 | PA14_72380 | *algB* | 6447033 | Δ10 bp |
| SM1073 | PA14_72380 | *algB* | 6447033 | Δ10 bp |
| SM1074 | PA14_10700 | *bphP* | 921631 | G → T |
| SM1149 | PA14_72380 | *algB* | 6446399 | Δ21 bp |
| SM1150 | PA14_10700 | *bphP* | 921151 | Δ12 bp |
| SM1151 | PA14_10700 | *bphP* | 921731 | G → T |

a: annotation from www.pseudomonas.com (Winsor et al., 2016)

**TABLE S3: Bacterial strains**

| Strain | Description | Reference |
|---|---|---|
| UCBPP-PA14 | Wild type *Pseudomonas aeruginosa* | Laboratory stock |
| SM32 | Δ*rhlR* | (Mukherjee et al., 2017) |
| SM1040 | Δ*rhlR* Δ*kinB* | This study |
| SM1050 | Δ*kinB* | This study |
| SM1111 | pUCP18-$P_{lac}$-*kinB* | This study |
| SM1112 | Δ*rhlR* Δ*kinB* pUCP18-$P_{lac}$-*kinB* | This study |
| SM1116 | Δ*kinB* pUCP18-$P_{lac}$-*kinB* | This study |
| SM1204 | *algB*$^{STOP}$ | This study |
| SM1212 | Δ*kinB algB*$^{STOP}$ | This study |
| SM1278 | Δ*kinB bphP*$^{STOP}$ | This study |
| SM1282 | *bphP*$^{STOP}$ | This study |
| SM1286 | pUCP18-$P_{lac}$-*algB* | This study |
| SM1303 | Δ*rhlR* pUCP18-$P_{lac}$-*algB* | This study |
| SM1326 | *bphP*$^{STOP}$ pUCP18-$P_{lac}$-*algB* | This study |
| SM1377 | *3xFLAG-algB* | This study |
| SM1378 | Δ*kinB 3xFLAG-algB* | This study |
| SM1386 | Δ*kinB bphP*$^{H513A}$ | This study |
| SM1387 | *algB*$^{STOP}$ pUCP18-$P_{lac}$-*algB* | This study |
| SM1388 | *bphP*$^{H513A}$ pUCP18-$P_{lac}$-*algB* | This study |
| SM1413 | *algB*$^{STOP}$ pBBR1-MCS5-$P_{lac}$-*3xFLAG-algB* | This study |
| SM1514 | *3xFLAG-algB kinB-SNAP* | This study |
| SM1523 | *3xFLAG-algB kinB*$^{P390S}$*-SNAP* | This study |
| SM1535 | pBBR-MCS5-$P_{lac}$-*bphP* | This study |
| SM1543 | *algB*$^{STOP}$ pBBR1-MCS5-$P_{lac}$-*bphP* | This study |
| SM1562 | *algB*$^{STOP}$ pBBR1-MCS5-$P_{lac}$-*3xFLAG-algB*$^{D59N}$ | This study |
| SM1563 | *algB*$^{STOP}$ pUCP18-$P_{lac}$-*algB*$^{D59N}$ | This study |
| SM1617 | *bphP-3xFLAG* | This study |
| SM1618 | *bphP*$^{H513A}$*-3xFLAG* | This study |

**TABLE S4: Plasmids**

| Plasmid | Description | Reference |
|---|---|---|
| pEXG2 | Allelic exchange vector with pBR origin, gentamicin resistance, *sacB* | (Hmelo et al., 2015) |
| pUCP18 | *E. coli-Pseudomonas* Amp$^r$ shuttle vector | Laboratory stock |
| pBBR1-MCS5 | *E. coli-Pseudomonas* Gent$^r$ shuttle vector | Laboratory stock |
| pIT2 | *ISlacZ/hah transposon mutagenesis vector* | (Jacobs et al., 2003) |
| pET21b | Protein expression vector, Amp$^r$ | Laboratory stock |
| pET28b | Protein expression vector, Kan$^r$ | Laboratory stock |
| pSP201 | pET21b-*bphP-His6* | This study |
| pSP202 | pET21b-*kinB-His6* | This study |
| pSP203 | pET28b-*His6-algB* | This study |
| pSP204 | pET21b-*bphP*$^{H513A}$*-His6* | This study |
| pSP205 | pET28b-*His6-algBPpu* | This study |
| pSP206 | pET28b-*His6-algB*$^{D59N}$ | This study |
| pSP207 | pET21b-*kinB*$^{P390S}$*-His6* | This study |
| pSP208 | pET28b-*His6-ntrC* | This study |
| pSP209 | pET28b-*His6-algBRce* | This study |
| pSP210 | pET28b-*His6-algBAxy* | This study |
